# Path-based reasoning for biomedical knowledge graphs with BioPathNet

**DOI:** 10.1101/2024.06.17.599219

**Authors:** Yue Hu, Svitlana Oleshko, Samuele Firmani, Zhaocheng Zhu, Hui Cheng, Maria Ulmer, Matthias Arnold, Maria Colomé-Tatché, Jian Tang, Sophie Xhonneux, Annalisa Marsico

## Abstract

Understanding complex interactions in biomedical networks is crucial for advancements in biomedicine, but traditional link prediction (LP) methods are limited in capturing this complexity. Representation-based learning techniques improve prediction accuracy by mapping nodes to low-dimensional embeddings, yet they often struggle with interpretability and scalability. We present BioPathNet, a novel graph neural network framework based on the Neural Bellman-Ford Network (NBFNet), addressing these limitations through path-based reasoning for LP in biomedical knowledge graphs. Unlike node-embedding frameworks, BioPathNet learns representations between node pairs by considering all relations along paths, enhancing prediction accuracy and interpretability. This allows visualization of influential paths and facilitates biological validation. BioPathNet leverages a background regulatory graph (BRG) for enhanced message passing and uses stringent negative sampling to improve precision. In evaluations across various LP tasks, such as gene function annotation, drug-disease indication, synthetic lethality, and lncRNA-mRNA interaction prediction, BioPathNet consistently outperformed shallow node embedding methods, relational graph neural networks and task-specific state-of-the-art methods, demonstrating robust performance and versatility. Our study predicts novel drug indications for diseases like acute lymphoblastic leukemia (ALL) and Alzheimer’s, validated by medical experts and clinical trials. We also identified new synthetic lethality gene pairs and regulatory interactions involving lncRNAs and target genes, confirmed through literature reviews. BioPathNet’s interpretability will enable researchers to trace prediction paths and gain molecular insights, making it a valuable tool for drug discovery, personalized medicine and biology in general.

## 1 Introduction

Biological entities interact in complex ways, crucial for sustaining life in living systems [1]. Understanding these interactions is central to systems biology, with network analysis playing a key role [2]. Biological networks are represented as graphs, where nodes can represent genes, proteins, diseases and more, and edges denote associations between them. Edges in a biological graph between genes can signify co-regulation or causal relationship (regulatory network) [3, 4], physical interactions (in protein-protein interaction networks (PPI) [5, 6]), as well as diseases-gene associations (like in disease-gene networks [7, 8]), among many.

Despite increasing high-throughput experiments, our grasp of biological networks is incomplete, leaving many interactions undiscovered. Due to the expense and time involved in wet lab experiments, computational methods such as link prediction (LP) are very important for inferring missing or potential associations within these networks based on the underlying topology [9]. LP is applied across network biology for diverse tasks ranging from predicting protein interactions over inferring gene regulatory networks to exploring pathways [10]. By revealing hidden connections, LP facilitates the discovery of biomarkers, drug targets, and insights into biological interactions [11, 12]. To predict potential relationships between unconnected nodes, one prevalent class of methods uses similarity metrics from traditional graph analysis, such as Personalized PageRank, Jaccard or Katz index [13, 14]. These metrics have been used for predicting disease-gene associations [15], including ncRNA-disease relationships and drug-disease associations [16].

While traditional graph metrics have been successful in biological link prediction, representation-based learning offers greater expressiveness for capturing the nuances and complexity of nodes in a graph. Nodes are mapped to low-dimensional vector representations called embeddings using shallow and deep non-linear transformations. Optimized embeddings position nodes with similar network neighborhoods closely in the embedding space so that links between nodes can be predicted based on their similarity in this space [17]. Methods include matrix factorization-based (e.g. Mashup [18]) and random walk-based approaches (e.g., DeepWalk [19], node2vec [20], struc2vec [21]). Network embedding techniques have found success in diverse domains, including drug repurposing, adverse drug reaction prediction, gene function prediction, and protein-protein interaction network completion, among others [22–25]. For example, Ruiz et al. [26] introduced the multiscale interactome, integrating disease-associated proteins, drug targets, and biological functions using biased random walks for node embeddings [26]. GeneWalk predicts gene functions via network representation learning with random walks [27]. Hu et al. [28] created a multi-modal network of genes and polygenic risk scores (PRS) for diseases, using DeepWalk for node embeddings to uncover associations between COVID-19 genes, co-morbidities, and genetic predispositions [28–30].

As opposed to the shallow learning approaches, methods such as Graph Convolutional Networks (GCNs) [31], Graph Autoencoders (GAEs) [32] and GraphSAGE [33] learn node embeddings from graph data using deep neural networks, by aggregating node messages from neighbors and learning a representation which reflects the neighborhood. Biological applications include OhmNet [24], which uses neural architectures to learn node embeddings in a multi-layer hierarchical network representing molecular interactions across human tissues, and Decagon, which [25] models polypharmaceutical side effects using GCNs and a multi-modal graph of protein-protein, drug-protein, and drug-drug interactions, enabling multi-relational link prediction with an encoder-decoder approach.

Early biological interaction models used basic networks, or uni-relational graphs, which failed to capture various entity associations’ semantics, such as distinguishing between inhibition and activation in protein-protein interactions. Recent efforts use heterogeneous multi-relational networks, or knowledge graphs (KGs), to better represent biological complexities by modeling facts as subject-predicate-object (SPO) triples. KG research is increasingly applied to tasks like question answering and information retrieval, with a key challenge being link prediction to complete KGs by estimating missing triplet components. Knowledge Graph Embedding (KGE) effectively learns low-rank representations of entities and relations, preserving graph structure and encoding relation semantics by optimizing a training loss that maximizes scores for positive triplets while minimizing those for corrupted triplets [34]. Representative KGE methods include TransE [34] for hierarchical relationships, DistMult [35] for symmetry patterns, ComplEx [36] for asymmetric relationships, and RotatE [37] for modeling symmetry, antisymmetry, inversion, and composition through rotational embeddings. A recent, expressive model that encodes indirect semantics using GNNs is the Relational Graph Convolutional Network (R-GCN) for multi-relational KGs [38]. R-GCN learns node embeddings by aggregating transformed feature vectors of neighboring nodes via a normalized sum and uses the DistMult factorization model for link prediction. Unlike conventional GCNs, R-GCNs introduce relation-specific transformations based on edge type and direction, making them suitable for multi-relational data in KGs. The study from Mohamed et al. [23] shows that KGE methods outperform traditional graph exploration methods in predicting drug-target interactions, polypharmacy side effects, and tissue-specific protein functions.

With the rapid accumulation of biomedical data, understanding disease biology and molecular factors’ roles in phenotypic outcomes is crucial for personalized diagnostics and treatments. KGs have become the dominant knowledge representation also in biomedicine, leveraging databases like UniProt [39], Gene Ontology [40, 41], and DrugBank [42]. LP tasks in biomedical KGs, such as Zhang et al.’s COVID-19 drug candidate exploration [43] with RotatE and DistMult, OntoProtein’s Gene Ontology-based KG for protein language model pretraining [44], and Biswas et al.’s node embedding algorithms for multi-modal biomedical KGs, enhance drug discovery and predict disease co-morbidities [45] via tensor factorization with complex-valued embeddings. Further, task-specific KGs and frameworks like BioCypher [46] further support KG construction, aiding predictive modeling for drug adverse reactions, repurposing, and biological concept associations.

While embedding-based approaches have shown significant performance in several benchmark tests, they are often limited to one-hop relations. In large biomedical KGs, relationships between entities are intricate and may involve multi-hop paths. Encoding a head entity without considering its specific tail entities requires embedding a vast amount of information (considering all possible tail entities). For large graphs, embedding all this information into a lower-dimensional vector is challenging and can lead to imprecise link predictions. Methods such as SEAL [47] and Grail [48] address the problem of predicting links between head and tail entities by embedding the subgraph structure around the link, encoding the two entities as a whole. However, these methods face scalability issues because they generate or materialize a subgraph for every link they try to predict. This process becomes a bottleneck when attempting to perform link prediction for all pairs.

To overcome these challenges, researchers started developing general and flexible representation learning frameworks for LP based on the paths between two nodes. The first application of this concept is the study from [49], who introduce KG4SL, a graph neural network (GNN) model that integrates KG message-passing for synthetic lethality (SL) prediction, leveraging a KG with 11 entity types and 24 relevant relationships associated with SL. Further, the Neural Bellman-Ford Network (NBFNet) introduces a novel framework for LP inspired by traditional path-based methods [50]. It represents node pairs as the sum of path representations, each derived from edge representations, and it employs a graph neural network with learned operators for efficient path formulation solutions, scalable to large graphs with low time complexity. NBFNet works with both homogeneous and multi-relational graphs, supporting LP across different graph types. Combining traditional path-based methods with GNNs, NBFNet demonstrates superior performance compared to node embedding methods. Additionally, path embedding methods offer better interpretability by visualizing important paths used for prediction, facilitating verification of biological plausibility.

To address link prediction in noisy biological KGs, we introduce BioPathNet. This message-passing neural network framework for path representation learning, inspired by NBFNet, specializes in predicting specific node subset relations within biomedical KG. As opposed to the node-embedding learning frameworks that optimize the embedding space based on one-hop relations, BioPathNet utilizes path-based reasoning to learn representations between source and target nodes based on relations along the path. BioPathNet makes use of a background regulatory graph (BRG), which may contain protein-protein interactions, as well as relationships between genes and other molecules with biomedical terms, being, therefore, more effective over prior path representation learning methods when it comes to predicting links on biomedical KGs. By leveraging additional graph information from the BRG for message passing, BioPathNet enriches path representations between node heads and tails, resulting in more precise predictions while avoiding learning irrelevant relationships. In addition, in BioPathNet, we introduce a stringent node type-aware negative sampling scheme that ensures contrastive learning and improves the decision boundary accuracy. These two points are especially important to large biomedical KGs that potentially encode noise derived from errors in experiments and, at the same time, are highly structured in how and which biological entities can interact.

We highlight BioPathNet’s effectiveness across four diverse LP tasks: *gene function prediction task*, *drug repurposing task*, i.e. disease-drug target interaction prediction in a zero-shot scenario, *synthetic lethality prediction task*, i.e. prediction of synthetic lethality gene pairs, *lncRNA-gene target prediction task*, i.e. inference of lncRNA-mRNA regulatory relationships. Despite varying KG requirements for each task, BioPathNet always surpasses KGE-based methods, including GNNs, in most of the tasks. For the *drug repurposing task* and *synthetic lethality prediction task*, it matches or outperforms task-specific models like TxGNN and KR4SL. BioPathNet discovers new drug-disease associations, including insights into Alzheimer’s disease, and scores potential lncRNA-mRNA interactions, validated against orthogonal datasets. Through examples, we demonstrate how BioPathNet enables the natural interpretation of predicted links, enhancing understanding of molecular disease mechanisms and regulatory processes.

## 2 Results

A knowledge graph (KG) is a heterogeneous directed graph comprising various types of entities (nodes) connected by relationships (edges). For instance, a KG might include nodes representing diseases, genes, and potential drug targets, with relationships such as ‘indication for’ or ‘involved in’ and model facts such as ‘drug A is an indication for disease B’ or ‘gene C is involved in disease D.’ KGs are typically represented as triples consisting of a head node, a tail node, and a relationship. The task of knowledge graph completion involves estimating the missing components of these triples. For example, one might predict the tail entities corresponding to a given head entity linked by a specific relationship, such as predicting diseases for which a particular drug is an indication, based on existing triples (i.e., existing knowledge). Knowledge graph completion methods can be broadly categorized into embedding-based and path-based approaches (Figure 1A). Embedding-based approaches use encoding models, ranging from simple linear models to complex neural networks, to learn feature representations of entities in a knowledge graph. These methods aim to preserve the structure of the original graph in a lower-dimensional space by minimizing the distance between the head and tail entity embeddings and the relationship embeddings or by maximizing the similarity between the embeddings of head entities, relations, and tail entities. Our path-based approach BioPathNet, on the other hand, can be leveraged to capture the structural information of KGs by learning representations for pairs of nodes (instead of single nodes) through paths. It learns node pair representations by parameterizing them as the generalized sum of path representations, with each path representation as the generalized product of edge representations along the path. (Figure 1B). This path formulation can be efficiently solved using the generalized Bellman-Ford algorithm based on dynamic programming. Moreover, the efficiency is further enhanced by learning the operators of the generalized Bellman-Ford algorithm with a message-passing graph neural network (see Methods).

**Fig. 1:**
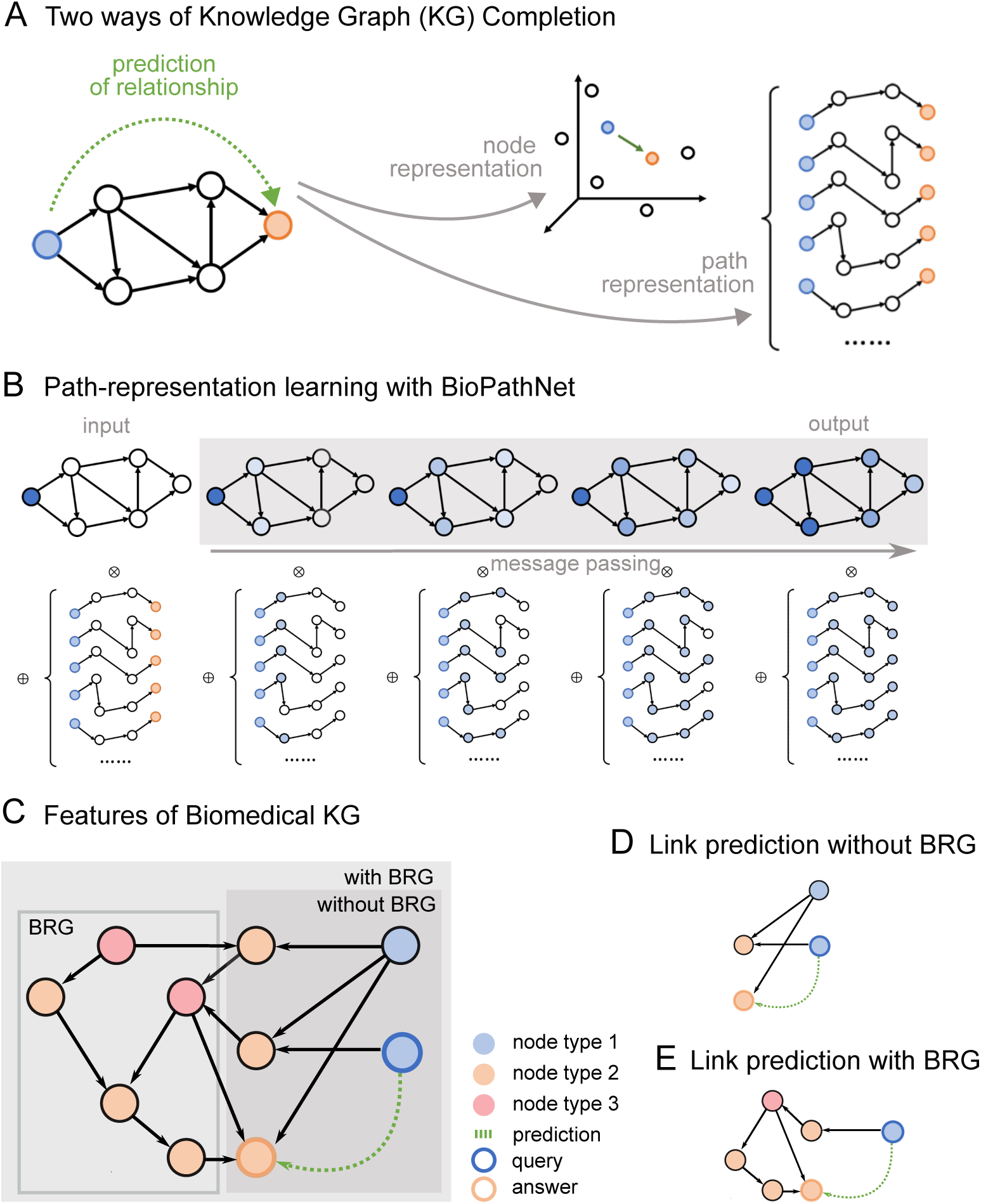
Link prediction (LP) in biological knowledge graphs: A) Inference of links using node-representation (node embedding) vs. path-representation learning. B) Illustration of the NBFNet framework, which uses the generalized Bellman-Ford algorithm to solve the shortest path problem between a head entity and tail entities via specific relationships, and employs message-passing GNNs to learn path representations, with a Multi-Layer Perceptron distinguishing positive and negative relationships. C) BioPathNet incorporates a background regulatory graph (BRG) to add additional gene connections, enhancing message passing and information flow beyond supervised training edges. It also uses an improved negative sampling scheme considering specific node types. D-E) Examples of prediction paths between head nodes (blue) and tail nodes (orange) in two scenarios are illustrated: D) a sub-graph without BRG, and E) a sub-graph that includes BRG connections used for learning. These examples also serve as model explanations, highlighting the paths that lead to the model’s predictions.

BioPathNet refines the NBFNet framework for biomedical KGs by using a stricter negative sampling strategy, where negatives are drawn from the same node type as positives, ensuring more challenging samples and better decision boundary learning. BioPathNet enhances prediction accuracy by integrating an external Biological Regulatory Graph (BRG) to improve entity connectivity during training’s message passing without affecting negative sampling and loss computation. Essentially, predictions can be made without and with a BRG, which is used solely for message passing (Figure 1C). For example, as illustrated in Figure 1D, when predicting the missing link between a head node and a tail node, messages can be passed between type 1 and type 2 nodes, resulting in a certain prediction path (Figure 1D). Alternatively, as illustrated in Figure 1E, a BRG can be integrated to further inform the predictions by leveraging additional knowledge bases, such as relations between type 2 and type 3 nodes. Besides enhancing performance, as demonstrated in the following sections, the incorporation of a BRG in BioPathNet allows the zooming into the molecular mechanisms behind a certain prediction. In fact, one can examine the sub-network (interaction partners, regulators) surrounding a specific node pair to derive a mechanistic hypothesis. This additional layer of insight leverages the broader biological context provided by the BRG.

To demonstrate BioPathNet’s versatility in performing graph completion across various tasks, we applied it to four link prediction challenges in biomedicine. These tasks vary in importance and difficulty, each involving heterogeneous KGs with distinct topological characteristics, sizes, and types of training data.

### Gene function prediction task

Our first goal was to evaluate the capacity, performance, and robustness of BioPathNet in biomedical KG link prediction, comparing its path embedding strategy to node embedding techniques, with a focus on the use of a BRG for message passing within the framework. For this, we conducted a proof-of-concept study focusing on gene function prediction. This involves assigning biological information, like terms corresponding to cellular pathways, to genes. Our approach involved applying BioPathNet to two scenarios: Firstly, we utilized a KG connecting genes and KEGG pathways through the relation ‘function of’ sourced from ConsensusPathDB [51], without a BRG. Secondly, we extended this KG by incorporating a BRG extracted from Pathway Commons [52–54] interactions encompassing genegene, chemical-gene, and chemical-chemical relationships. The objective of this experiment was two-fold: firstly, to evaluate BioPathNet’s performance in link prediction compared to traditional node embedding methods, and secondly, to assess the impact of augmenting the KG with a BRG on enhancing the accuracy of gene function annotation tasks. Through this investigation, we aimed to validate BioPathNet’s utility in leveraging complex biomedical data structures for improving predictive modeling in gene function annotation within KGs.

In direct comparison with Knowledge Graph Embedding (KGE)-based methods such as TransE, DistMult, and RotatE, as well as Graph Convolutional Networks (R-GCN), BioPathNet demonstrated consistently superior performance across different metrics (Figure 2B-C). In the setting without utilizing the BRG, BioPathNet achieved a Mean Reciprocal Rank (MRR), which measures how well the model ranks the correct pairs, of 0.464, outperforming the KGE methods, which averaged 0.371, and R-GCN, which achieved 0.348. For the Hits@k metric, which indicates the percentage of ground truth items captured within the top *k* predictions, BioPathNet obtained 63.5% in the top 10 predictions, compared to RotatE’s 56.5% (Figure 2B). Upon leveraging the BRG for biological regulation-enhanced message passing, performance improvements were observed primarily for R-GCN and BioPathNet. R-GCN’s MRR increased marginally from 0.348 to 0.355, whereas BioPathNet’s performance rose from 0.464 to 0.549, corresponding to an 8.5% gain. In terms of capturing ground truth positives within the top 10 predictions, BioPathNet excelled with 72.6%, whereas R-GCN achieved 53.1% (Figure 2C). Interestingly, KGE methods did not leverage the additional BRG information effectively; in fact, the TransE model’s performance dropped significantly from 0.376 to 0.272, indicating a disadvantage rather than an enhancement in predictive capability. By conducting experiments for each method over 5 different model seeds, we observed standard deviation for each method ranging between 0.01 and 0.03, yielding robust predictions for all methods (Figure 2B).

**Fig. 2:**
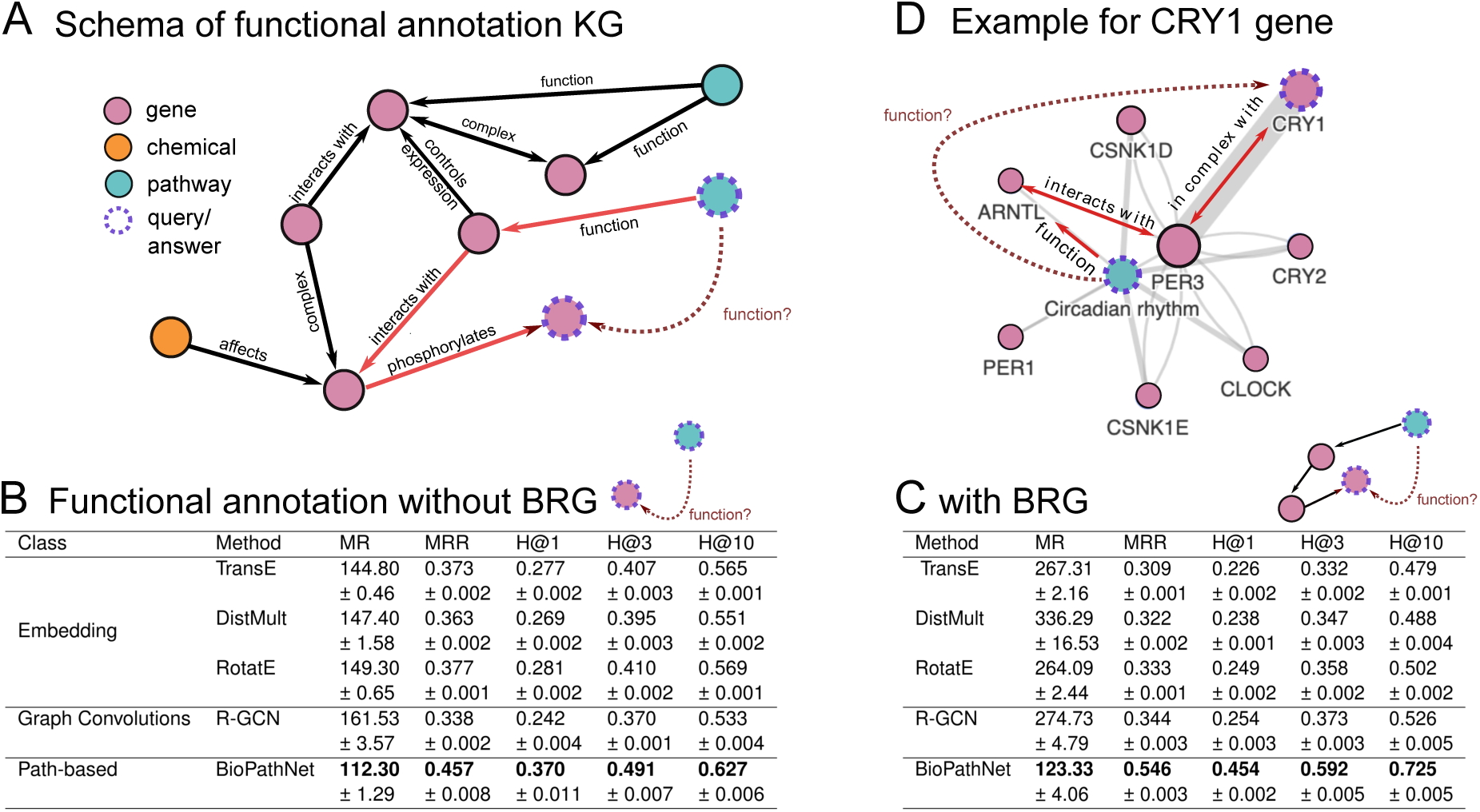
Benchmark of knowledge graph completion algorithms on the gene function annotation task: A) Illustration of BioPathNet leveraging a BRG encompassing genes, chemicals, and cellular pathways, to predict gene functions, i.e. associations of genes with specific cellular pathways. B-C) Performance on the gene function prediction task against classical KGE-based methods, namely TransE, DistMult, RotatE and R-GCN for link prediction. B) without the underlying BRG and C) with the BRG. Metrics reported for comparison are mean rank (MR), mean reciprocal rank (MRR), and Hits at 1, 3, and 10. D) The visualization highlights the significant paths employed by the BioPathNet model to predict a link between *CRY1* and the *Circadian rhythm*. The top 10 paths are depicted, where the width of each edge corresponds to its weight, and the path with the highest weight is highlighted in red.

One key advantage of NBFNet is its ability to provide interpretable predictions through paths, which are crucial for understanding the rationale behind specific predictions. Intuitively, these interpretations should highlight paths that significantly influence the prediction. In BioPathNet, this follows the NBFNet framework (see Methods), where the top-*k* path interpretations for a prediction are formally defined as the first derivative (gradient of the prediction) with respect to each path between a head and a tail node. In this task, we show an example of how BioPathNet interprets its predictions and visually presents the most critical paths for predicting the function of the *CRY1* gene, specifically its association with the *Circadian rhythm* pathway (Figure 2D). The figure illustrates the top 10 most significant paths ranked by gradient, where the width of each edge reflects how frequently it appears among these top paths. Additionally, the most crucial path, ranked highest by gradient, is highlighted in red, encompassing: *CRY1* in complex with → *PER3* interacts with → *ARNTL* before feeding into the pathway *Circadian rhythm* over the relation function of (Figure 2D). The retrieved path makes sense as it recovers the well-known mechanisms by which the essential transcription factors controlling the cellular circadian rhythm, ARNTL, and CLOCK, upregulate the expression of PER3 and CRY2 [55, 56]. They, in turn, form heterodimers to repress their own expression, creating a negative feedback loop of regulation [57, 58].

### Drug repurposing task

In the second part of our study, we evaluated BioPathNet in a more challenging scenario: predicting new drug candidates for diseases by repurposing existing drugs indicated for other conditions. This drug repurposing task was conducted in a zero-shot scenario, where the target disease has minimal molecular characterization and no available treatments. For this experiment, we followed the data split procedure implemented in TxGNN [59], a state-of-the-art graph neural network model designed to predict drug-disease relationships in zero-shot scenarios, which builds embeddings of nodes and relations from a comprehensive biomedical knowledge graph, the PrimeKG knowledge graph (Supplementary Figure 1A) [60] (see Methods for more details).

More in detail, TxGNN creates 5 ‘disease area’ splits to simulate zero-shot conditions, ensuring that diseases in the test set used for inference (1) have no approved drugs in the training data, (2) have limited overlap with the training diseases by excluding similar ones, and (3) lack molecular data by removing their biological neighbors from the training set. These splits provide challenging yet realistic evaluation scenarios, mimicking zero-shot drug repurposing (see Methods). These splits create challenging yet realistic evaluation scenarios for zero-shot drug repurposing by simulating a new disease with minimal knowledge, no similar diseases, and no known treatments. Connections to treatments and most biological neighbors are removed from the training set to prevent their use in message passing. Five distinct zero-shot disease areas were used: adrenal gland, anemia, cardiovascular, cell proliferation, and mental health.

The BioPathNet model used approximately 5.7 million directed edges solely for message passing in each prediction setting (Supplementary Table 4), including non-drug-disease edges like protein-protein and disease-disease relations. In contrast, edges used for both message passing and supervision were limited to drug-disease interactions, such as ‘contraindication’ and ‘indication’. On average, the training set contained around 33,000 edges, and the validation set around 4,000. The number of testing edges varied significantly between disease areas, with 1,047 contraindications and 999 indications in the cell proliferation split, compared to 303 contraindications and 33 indications in the adrenal gland disease area (Supplementary Table 4).

For each disease area split, we evaluated BioPathNet against TxGNN by assessing their performance in predicting ground truth drugs for the relations ‘contraindication’ and ‘indication’. Specifically, we ranked all drugs (tail node) based on their likelihood of being an indication or contraindication for a specific disease (head node). This involved computing the probability for each drug to be an indication or contraindication for a disease, *p*(drug | disease, relation), from both BioPathNet and TxGNN (Figure 3A). In the comparison, BioPathNet achieved higher AUPRC than TxGNN in two out of five disease areas for contraindication prediction and in all disease areas for indication prediction (Supplementary Figure 1B). The difference in performance, Δ, is calculated by subtracting TxGNN’s AUPRC from BioPathNet’s AUPRC; thus, a positive Δ indicates better performance by BioPathNet. For contraindications, TxGNN outperformed BioPathNet in adrenal gland, cardiovascular, and mental health areas with Δ values of −4.6, −0.2, and −2.0 percentage points, respectively. Conversely, BioPathNet outperformed TxGNN in anemia and cell proliferation with Δ values of 4.8 and 9.0. In the indication prediction task, BioPathNet consistently had positive Δ values, ranging from 5.9 to 22.6 percentage points (Figure 3B). To summarize the performance in a single metric, the AUPRC was averaged across contraindications and indications for each disease area split (Supplementary Table 7). For cell proliferation, the difference in performance Δ was 0.119 (0.556 − 0.437), representing a performance increase for BioPathNet over TxGNN of 27.3%. Similarly, the increases were 17.1%, 14.1%, 25.9%, and 16.9% for adrenal gland, anemia, cardiovascular, and mental health, respectively. On average, BioPathNet outperformed TxGNN by 20.2% across all disease area splits.

**Fig. 3:**
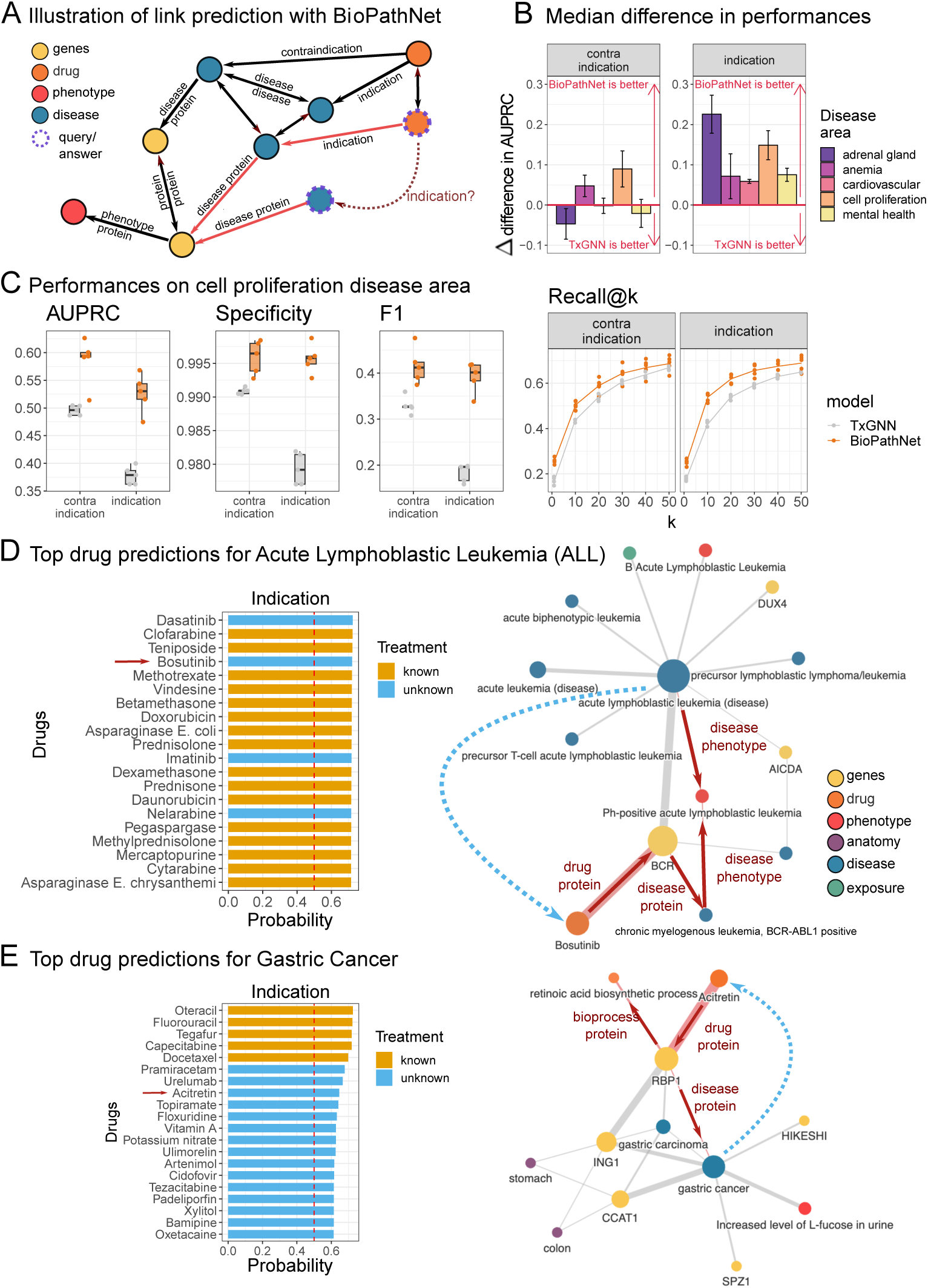
Comparison of BioPathNet and TxGNN model on the drug-disease relations prediction task: A) Schematic of the PrimeKG graph used by BioPathNet and illustration of a drug-disease indication relationship. B) Mean AUPRC differences between BioPathNet and TxGNN across five disease area splits (adrenal gland, anemia, cardiovascular, cell proliferation, mental health). A positive delta indicates higher AUPRC for BioPathNet. C) Performance metrics for the cell proliferation split. Recall@k reflects the proportion of ground truth edges in the top k predictions, reported for contraindication and indication. D) Acute Lymphoblastic Leukemia and E) Gastric Cancer within the cell proliferation area. Left panels show predicted drug indications for ALL (D) and Gastric Cancer (E), ranked by BioPathNet prediction probability. Known indications are orange; novel indications are light blue. The right panels visualize the gradient importance of paths predicting Bosutinib for ALL (D) and Acitretin for Gastric Cancer (E), showing the top 10 significant paths with edge widths representing weights and the highest-weight path in red.

#### Cell Proliferation Split

A detailed breakdown of BioPathNet and TxGNN in terms of Specificity, F1, and Recall@k for the *cell proliferation* disease area split is illustrated in Figure 3C. Both models performed well in identifying true negatives, with BioPathNet showing only slightly higher specificity (0.996 vs. 0.981) in the indication setting. The F1 score, representing the balance between precision and sensitivity, was 0.415 for BioPathNet vs. 0.330 for TxGNN in the contraindication setting, and 0.393 vs. 0.183 for indication. This trend, observed in the cell proliferation split, holds across all other disease splits. Notably, BioPathNet showed a greater improvement in performance for indication than contraindication, though with slightly higher variance compared to TxGNN. In the cell proliferation split, 178 diseases had known indications, with an average of 5.58 indications per disease. For 60 out of 178 diseases, all known treatments were prioritized within the top 10 predictions sampled from a list of 7,000 drug candidates. Recall@k quantifies the proportion of ground truth items found within the top *k* predictions. For instance, at *k* = 20, the recall for indication across all diseases was 0.619 for BioPathNet, meaning that 61.9% of the ground truth drugs were found within the top 20 predictions. In contrast, Recall@20 for TxGNN was 0.539.

### Case study from Cell Proliferation: Acute Lymphoblastic Leukemia (ALL)

After quantitatively evaluating the performances, we further examined individual disease predictions within the Cell Proliferation split. Among the best-performing models was Acute Lymphoblastic Leukemia (ALL), which is a complex cancer involving abnormal proliferation of lymphoid cells in blood and bone marrow, impairing immune function [61]. Commonly observed chromosomal aberrations include the t(9;22) translocation, which produces the constitutively active tyrosine kinase BCR-ABL1, associated with Philadelphia chromosome-positive ALL [62].

We used BioPathNet for the prediction of the drugs associated with ALL. We were able to correctly predict the only known contraindication - drug Aprostadil on rank 1 with a probability score of 0.727, as well as all 21 known indications within the top 34 predictions (Figure 3D). Upon investigating the top indication predictions, we identified the highest-ranked known treatments (in orange) Clofarabine, Teniposide, and Methotrexate. Additionally, the top-ranked unknown treatments (in blue) were Dasatinib and Bosutinib (Figure 3D, left). We further set out to interpret our predictions by visualizing the most important paths for the predictions. The visualization plot summarizes the top ten most important paths as ranked by gradient (see Methods), with the edge width reflecting the number of times the edge appears among the top ten paths. The first novel indication prediction with a probability score of 0.724 was Dasatinib (Figure 3d, left), which is not present in the ground truth database PrimeKG. However, Dasatinib, an inhibitor of the constitutively active tyrosine kinase BCR-ABL, is already used for treating Philadelphia chromosome-positive acute lymphoblastic leukemia (Ph+ ALL) in cases of resistance or intolerance to prior therapies. The next novel prediction, Bosutinib, is an unknown drug predicted to treat ALL with a probability score of 0.721 (Figure 3D, left). To gain confidence in this prediction, we visualized the most important paths leading to it, focusing on the local subgraph to explain our results. For Bosutinib as an indication for ALL, the similarity to other (lymphoblastic) leukemia types was revealed, along with significant disease genes AICDA and DUX4 [63–65]. The most crucial path passes through the phenotype *Ph+ ALL*, the disease *chronic myelogenous leukemia, BCR-ABL1 positive*, and the gene *BCR*, before connecting to *Bosutinib* via the *drug protein* relation (Figure 3D, right). Indeed, Bosutinib was originally indicated for chronic myeloid leukemia in 2012 [66, 67] and is currently being investigated for the treatment of ALL [68].

#### Hypothesis generation for treatment of Gastric Cancer

To demonstrate BioPathNet’s ability to generate hypotheses for lab testing and evaluation, we investigated gastric cancer. Similar to ALL, both known contraindications and indications were ranked highly (all 5 contraindications in the top 6, and 5 out of 6 indications in the top 5) (Figure 3E, left). One novel drug predicted for gastric cancer treatment was Acitretin, an oral retinoid similar to Vitamin A, indicated for skin diseases like psoriasis by inhibiting excessive cell growth and keratinization [67]. Although untested for gastric cancer, Acitretin has been considered in combination with Clarithromycin for cutaneous squamous cell carcinoma due to its apoptosis-inducing properties [69, 70]. Interestingly, all paths from gastric cancer to Acitretin in the interpretability plot pass through RBP1 (retinol-binding protein 1), annotated with the *retinoic acid biosynthesis process* (Figure 3E, right). Recent studies suggest this pathway’s involvement in gastric cancer treatment and provide pre-clinical evidence supporting the use of All-Trans Retinoic-acid (ATRA) [71, 72]. By visualizing important paths between drugs and diseases, researchers can verify predictions’ plausibility and generate hypotheses for further laboratory validation.

#### Predicting drug indications for Alzheimer

For the final experiment in the drug repurposing task, we aimed to investigate a disease not analyzed by TxGNN to evaluate how well BioPathNet generalizes to a novel case study. Hereby, we examined the indication predictions for Alzheimer’s disease (AD) together with medical experts. AD is a neurodegenerative disorder characterized by extracellular amyloid beta and intracellular tau protein accumulation in the brain. These neuropathological changes occur decades before clinical symptoms, ultimately leading to synapse loss, brain atrophy, and dementia symptoms like memory loss and behavioral changes. While amyloid and tau are central to AD, the exact mechanisms remain unclear. Emerging evidence suggests additional pathways, such as immunoinflammation and bioenergetic dysregulation, may offer promising therapeutic targets [73–75]. Presently, FDA-approved treatments include only two disease-modifying and five symptomatic treatments, none of which provide a cure for AD. To explore the potential of BioPathNet for such complex and heterogeneous diseases, we trained BioPathNet on a data split tailored for zero-shot prediction on a custom-defined Alzheimer’s disease area split. Here, we followed the disease evaluation code as provided by TxGNN to exclude all treatments for Alzheimer’s, as well as closely related diseases (e.g. dementia)(Supplementary Table 5). We then evaluated the top 20 predictions for indications and contraindications for Alzheimer’s disease.

Among the top 14 predictions, seven out of eight drugs classified as known treatments according to PrimeKG, and four out of seven FDA-approved treatments for Alzheimer’s disease (AD), were successfully retrieved (Figure 4A). Additionally, the model identified Epicriptine, a nootropic drug with an unknown mode of action, and Acetylcarnitine, which is functionally involved in *β*-oxidation of fatty acids [76]. Known AD drugs, which obtained a low probability from BioPathNet were Pramiracetam (ranked 344), used for cognitive impairment in aging and dementia [77], and FDA-approved treatments such as Memantine (ranked 412), an *N* -methyl-D-aspartate receptor antagonist [78], the recently approved monoclonal antibodies Lecanemab (ranked 2250), and retracted Aducanumab (ranked 2216) [79, 80].

**Fig. 4:**
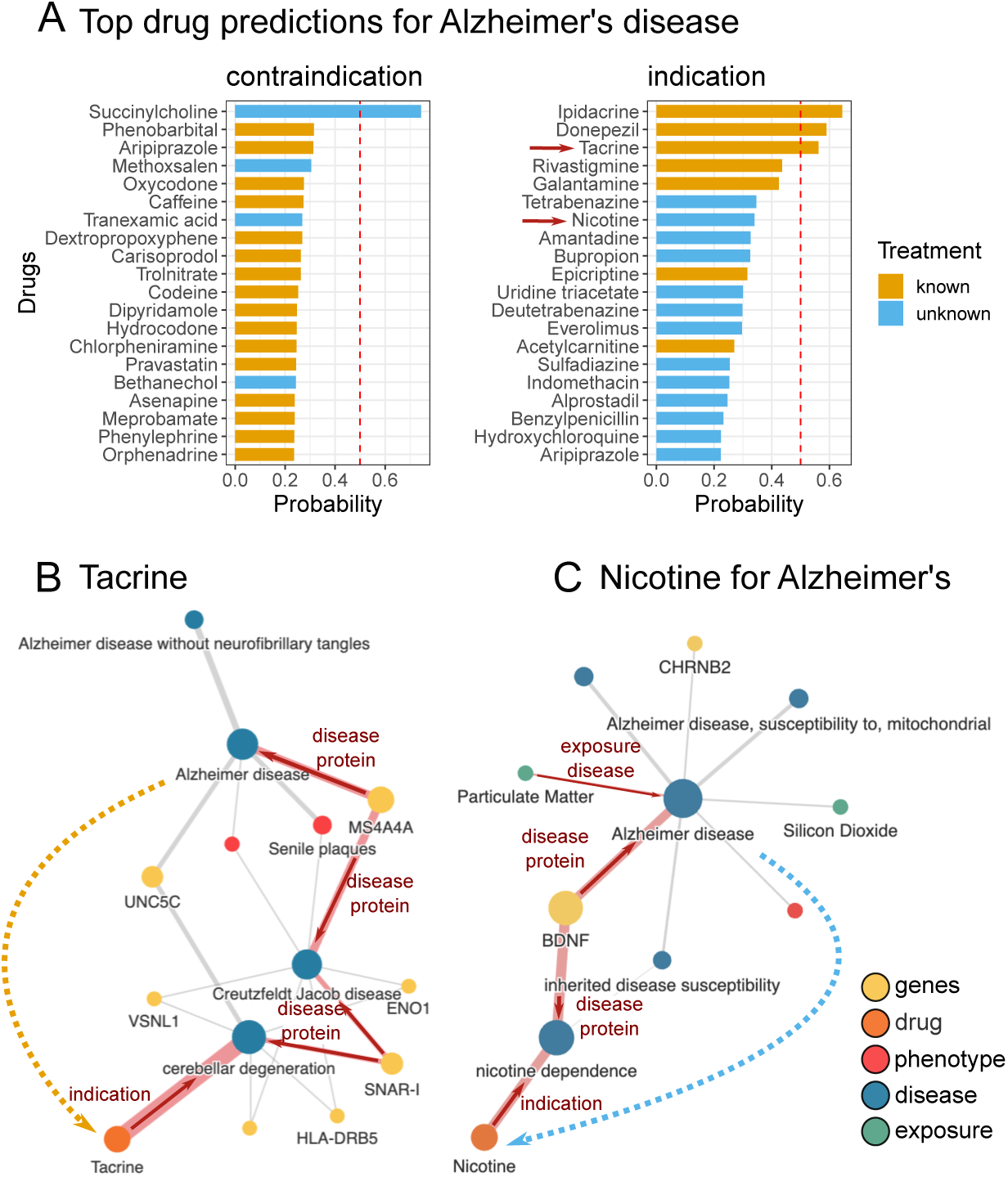
Predictions of BioPathNet on custom data split of Alzheimer’s disease: A) Top 20 predictions of contraindication and indication for custom Alzheimer’s disease, ranked by BioPathNet prediction probability. Known treatments, included in the ground truth of PrimeKG, are highlighted in orange, while novel indications are in light blue. Visualization of gradients on path importance for the prediction of B) Tacrine (a known treatment for Alzheimer) and C) Nicotine for Alzheimer’s (a newly predicted indication). The visualization shows the top 10 significant paths used by BioPathNet for prediction, with edge widths representing weights and the highest-weight path highlighted in red.

Interestingly, two drugs currently undergoing clinical trials were among the top 20 predicted novel indications: Nicotine, a nicotinic acetylcholine receptor agonist which is being tested in a Phase II clinical trial (NCT02720445) to improve cognition, and Bupropion, an *N* -methyl-D-aspartate receptor antagonist that is being tested as a component of the drug AXS-05 in two Phase III (NCT05557409, NCT04947553) clinical trails [81] to help with agitation associated with AD. Examination of the interpretability graphs shows that both predictions are associated with the brain-derived neurotrophic factor (BDNF), a gene crucial for synaptic maintenance and plasticity in the brain [82] (Figure 4C). Synaptic plasticity plays a pivotal role in AD [83], with research indicating lower levels of BDNF in both blood [84] and brain [85] in AD patients and linking higher levels of brain BDNF with slower cognitive decline [86] in elderly individuals. Both predicted drugs, Bupropion and Nicotine, have demonstrated an ability to elevate BDNF levels in serum [87, 88], providing a functional hypothesis for the mechanism of these drugs in the context of AD. Another promising candidate predicted with high probability was Everolimus, an analog of Rapamycin and a selective inhibitor of the mammalian target of Rapamycin (mTOR) kinase signaling pathway. This pathway has been implicated in both normal aging and pathological aging processes, making it a promising target for intervention, particularly in the early stages of disease onset [89]. Currently, Rapamycin is being evaluated as a potential disease-modifying therapy in Phase II (NCT04629495) and Phase I (NCT04200911) clinical trials involving older adults with mild cognitive impairment or early AD [81]. Furthermore, Rapamycin has shown beneficial effects on amyloid and tau burden in mouse models of AD [90]. Everolimus, although structurally similar to Rapamycin, has favorable clinical pharmacokinetics that influence, for example, bio-availability and tissue distribution [91]. Therefore, Everolimus may present an additional candidate for targeting the hyperactivated mTOR pathway in AD.

### Synthetic lethality prediction task

SL occurs when the simultaneous mutation of two genes leads to cell death, while the mutation of either gene alone is non-lethal [92]. We next examined the prediction of missing SL gene pairs with BioPathNet, which is of high interest in anti-cancer drug treatment. In fact, when the first partner of a gene pair is inhibited by mutations in cancer cells, targeting the second partner can induce selective cell death in cancer cells without harming normal cells. This approach is crucial when direct targeting of cancer driver genes is impractical, but their SL partners offer viable treatment alternatives. Given the great potential to design personalized treatments through SL-based therapy, computational methods to predict novel gene interaction partners are of great importance. In our study, we leverage BioPathNet for this task and compare it against the state-of-the-art method, named KR4SL [49], which is a path-representation learning GNN-based method to predict and explain synthetic lethality gene pairs (see Methods and Extended Methods section in the Supplementary File).

For training and inference of BioPathNet, we used the SynLethDB-v2.0 [93] data pre-processed by the authors of KR4SL (Supplementary Table 8). SynLethDB is a database that compiles SL pairs from biochemical assays, related databases, computational predictions, and text mining. Each SL relation in the database is assigned an integrative confidence score, prioritizing experimental evidence and giving higher scores to pairs supported by multiple sources.

To enhance model training with reliable SL pairs, we focused on those primarily from experiments and partially from computational predictions and text mining (Supplementary Figure 3A). We excluded pairs with confidence scores below 0.3, removing over 25% of computational and text-mined pairs. Specifically, 3,138 of 9,327 computationally predicted pairs and 905 of 5,614 text-mined pairs were discarded. This resulted in a training set of 8,770 SL pairs, a validation set of 3,172, a test set of 6,254, and a known SL set in BRG of 13,161 pairs (Supplementary Table 9). The filtered set is referred to as ‘thresholded data,’ while the original is ‘unthresholded data.’ Setting thresholds below 0.3 was impractical: a threshold of 0.1 removed no SL pairs, while 0.2 removed less than 10% of non-experimental pairs. We tested thresholds from 0 to 0.8 in 0.1 increments to evaluate their impact on the performance of BioPathNet and KR4SL (Supplementary Figure 3B).

For each seed and threshold, KR4SL and BioPathNet were evaluated on NDCG@k, Precision@k, and Recall@k for *k* ∈ 10, 20, 50 (Supplementary Figure 3B). BioPathNet significantly outperformed KR4SL in unthresholded data (p-value *<* 0.01, one-sided t-test, Figure 5B) and in thresholded data (p-value *<* 0.1, one-sided t-test) for Recall@10 and Precision@10, as well as for other metrics (p-value *<* 0.01, one-sided t-test, Figure 5B). For threshold 0.2, BioPathNet also signifcantly outperformed KR4SL for Precision@10 (p-value *<* 0.05, one-sided t-test), as well as for other metrics (p-value *<* 0.01, one-sided t-test, Supplementary Figure 3C). With a threshold of 0.3, BioPathNet achieved the best overall performance for all metrics (Supplementary Figure 3B). Although higher thresholds (0.4, 0.5, 0.6) showed improved performance, we chose a threshold of 0.3 for BioPathNet to balance training data quality and variance for more reliable predictions.

**Fig. 5:**
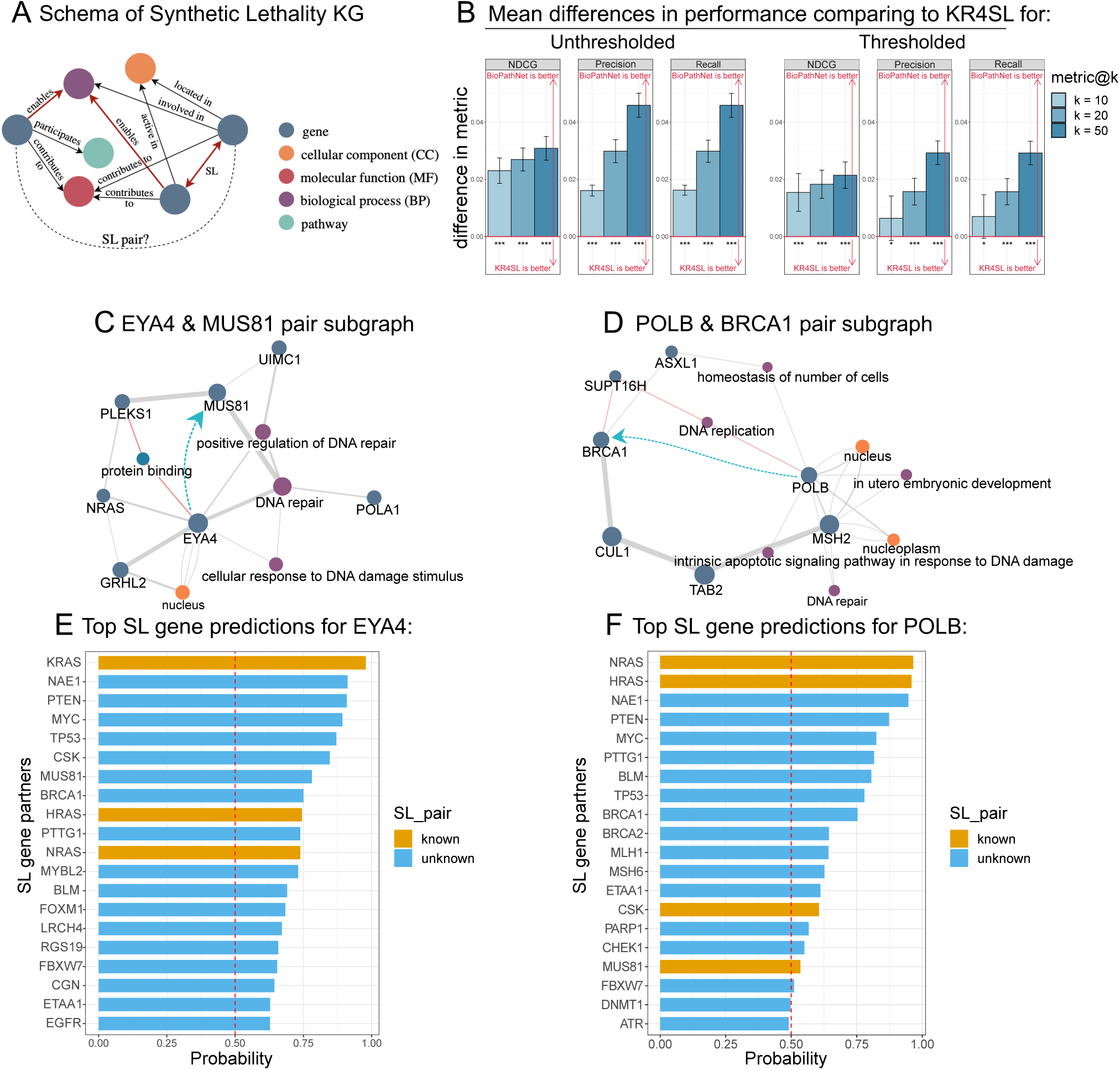
Comparison of BioPathNet with state-of-the-art SL gene pair prediction algorithm KR4SL: A) Illustration of SynLeth KG for the prediction of SL gene pairs, consisting of genes and their SL interactions, cellular component (CC), molecular function (MF), biological process (BP) and pathway. B) Mean difference in performances between BioPathNet and KR4SL given as NDCG, Precision, and Recall for both unthresholded and thresholded data. C) Visualization of gradients on paths important for the prediction of the EYA4 - MUS81 pair. D) Visualization of gradients on paths important for the prediction of the POLB - BRCA1 pair. E) Top predicted SL gene partners for EYA4. F) Top predicted SL gene partners for POLB.

A detailed breakdown of the performance of both methods in terms of MRR, NDCG@k, Precision@k, and Recall@k for *k* ∈ {10, 20, 50} is reported for each model run in Supplementary Tables 12 and 13 for unthresholded and thresholded data, respectively.

After evaluating model performance, we assessed BioPathNet’s ability to identify novel SL gene pairs using thresholded data, focusing on new, consistently predicted SL partners across model runs, predicted with an average MRR above 0.75. We analyzed the novel gene pair EYA4 and MUS81, where EYA4, involved in transcription, eye development, and DNA repair, is linked to hearing loss and cardiomyopathy, while MUS81 is essential for DNA repair. BioPathNet ranks MUS81 as the 7th synthetic lethality partner for EYA4 (Figure 5E). Figure 5C shows the explanation subgraph with multiple paths from EYA4 to MUS81 through shared processes like DNA repair, supporting their SL relationship.

Another example involved POLB and BRCA1. POLB, a repair polymerase essential for base-excision repair and linked to Werner syndrome and esophageal cancer, is consistently predicted across seeds as SL partner for POLB among the top 20 candidates (Figure 5F). The explanation subgraph in Figure 5D shows the top 10 paths from POLB to BRCA1, highlighting shared biological processes such as DNA repair, DNA replication, cellular homeostasis, and apoptotic signaling. Notably, POLB and MSH2 share nodes related to DNA repair, while POLB and SUPT16H (an SL partner of BRCA1) are involved in DNA replication. Additionally, POLB and ASXL1 (another known SL partner of BRCA1) share cellular homeostasis, supporting the evidence for the SL relationship between POLB and BRCA1.

### LncRNA-target prediction task

Long non-coding RNAs (lncRNAs) are a heterogeneous group of transcripts that lack protein-coding potential, usually longer than 200 nt. They encompass a substantial portion of the genomes of complex organisms. The extensive transcription of these non-coding transcripts unveils a significant shift in our understanding of the pivotal role of RNAs in gene regulation [94]. LncRNAs play crucial roles in imprinting control, immune response, epigenetic regulation, and gene regulatory networks. Their mutations and dysregulation are linked to numerous diseases, making them valuable biomarkers for diagnosis, treatment, and prognosis. Data from consortia like ENCODE [95] and FANTOM5 [96], along with resources such as RNAcentral [97] and NONCODE [98], estimate over 200,000 potential lncRNA transcripts, highlighting their diverse functional roles and mechanisms. Long non-coding RNAs regulate gene expression both locally (cis) and distantly (trans) by interacting with RNA Binding Proteins (RBPs) and other nucleic acids. They can function as signals, scaffolds, guides, and enhancer-like RNAs, modulating gene expression through chromatin looping, recruiting repressive complexes, like in the case of XIST and HOTAIR or competing endogenous RNAs (ceRNAs) in the cytoplasm, where they can act as microRNA sponges or decoys. Despite recent advances, most lncRNAs remain functionally uncharacterized, and their roles in disease biogenesis and progression are still unknown.

The imperative task of elucidating the functions and mechanisms of numerous lncRNAs underscores the urgency of identifying their targets using both experimental and computational approaches, which is a crucial initial step in functional analysis. Identifying the targets of lncRNAs, whether proteins, RNA sequences, or chromatin, is crucial in lncRNA research. Experimental methods like RNA pull-down, ChIRP, RIP, and CLIP systematically screen and identify lncRNA targets, enabling the construction of regulatory networks.

On the KG derived from the lncRNA regulatory graph in LncTarD 2.0 (Figure 6A), BioPathNet significantly outperformed both node embedding-based methods and the basic NBFNet algorithm across all metrics (Table 1). The lower performance of embedding-based methods highlights the importance of considering node and gene types in gene regulatory knowledge graph completion, an aspect neglected by the basic versions of TransE and DistMult used in this study. Additionally, our experiments demonstrate the effectiveness of BioPathNet’s negative sampling strategy and the successful integration of the external BRG (Table 1). To demonstrate BioPathNet’s ability to uncover novel lncRNA regulations, we focused on the lncRNA PVT1, a Myc regulator frequently over-expressed in cancers, crucial for tumor initiation, proliferation, invasion, and apoptosis, and linked to poor prognosis and therapy resistance. Using a trained model, we performed link prediction with PVT1 as the viewpoint, i.e. we set PVT1 as the head node and computed all conditional probabilities *p*(*t*|PVT1*, r*) for all nodes across all relationship types. The top 5 novel predictions with the highest probabilities are reported in Figure 6B and Table 2, where “novel” indicates the absence of a direct connection between these genes in the knowledge graph. Additionally, the top 10 most crucial paths for these predictions, ranked by gradient, are illustrated in Figure 6 and Supplementary Figure 3. For all five predictions involving PVT1, the model identified significant edges that do not form a direct path from PVT1 to the target gene. Instead, the predictions are inferred through bipartite graphs, where PVT1 and other lncRNAs are on one side, and the PVT1 target gene, along with other co-regulated genes, are on the other. This aligns with the biological understanding that genes in the same cluster are often regulated by the same factors. For example, Figures 6C and 6D show that PVT1 and GIHCG interact with the EZH2 protein, while GIHCG epigenetically regulates MIR200A, MIR200B, and MIR429. Thus, the model infers that MIR200A and MIR429 are also regulated by PVT1. This prediction is meaningful because it is known that the MIR200 family, which includes MIR200A, MIR200B, MIR429, MIR141, and MIR200C, is crucial in cancer initiation and metastasis. Evidence in the literature also indicates that PVT1 promotes cervical cancer progression by silencing MIR200B through EZH2 interaction, leading to histone H3K27 trimethylation and MIR200B inhibition [99]. PVT1 may also influence melanoma by regulating MIR200C via EZH2 [100]. Notably, PVT1-EZH2 regulation appears in all five novel predictions (Figure 6 and Supplementary Figure 4), underscoring EZH2’s role in PVT1 regulation. This aligns with experimental evidence of PVT1-EZH2 interactions in various cancers, including gastric, thyroid, glioma, and hepatocellular carcinoma. Furthermore, BioPathNet predicts an interaction between PVT1 and SUZ12 (Supplementary Figure 4), a member of the Polycomb Repressive Complex 2 (PRC2), along with EZH2. The model identifies a path through the physical interaction of the oncogenic lncRNA APTR, which represses the CDKN1A/p21 gene promoter via PRC2, involving both EZH2 and SUZ12.

**Fig. 6:**
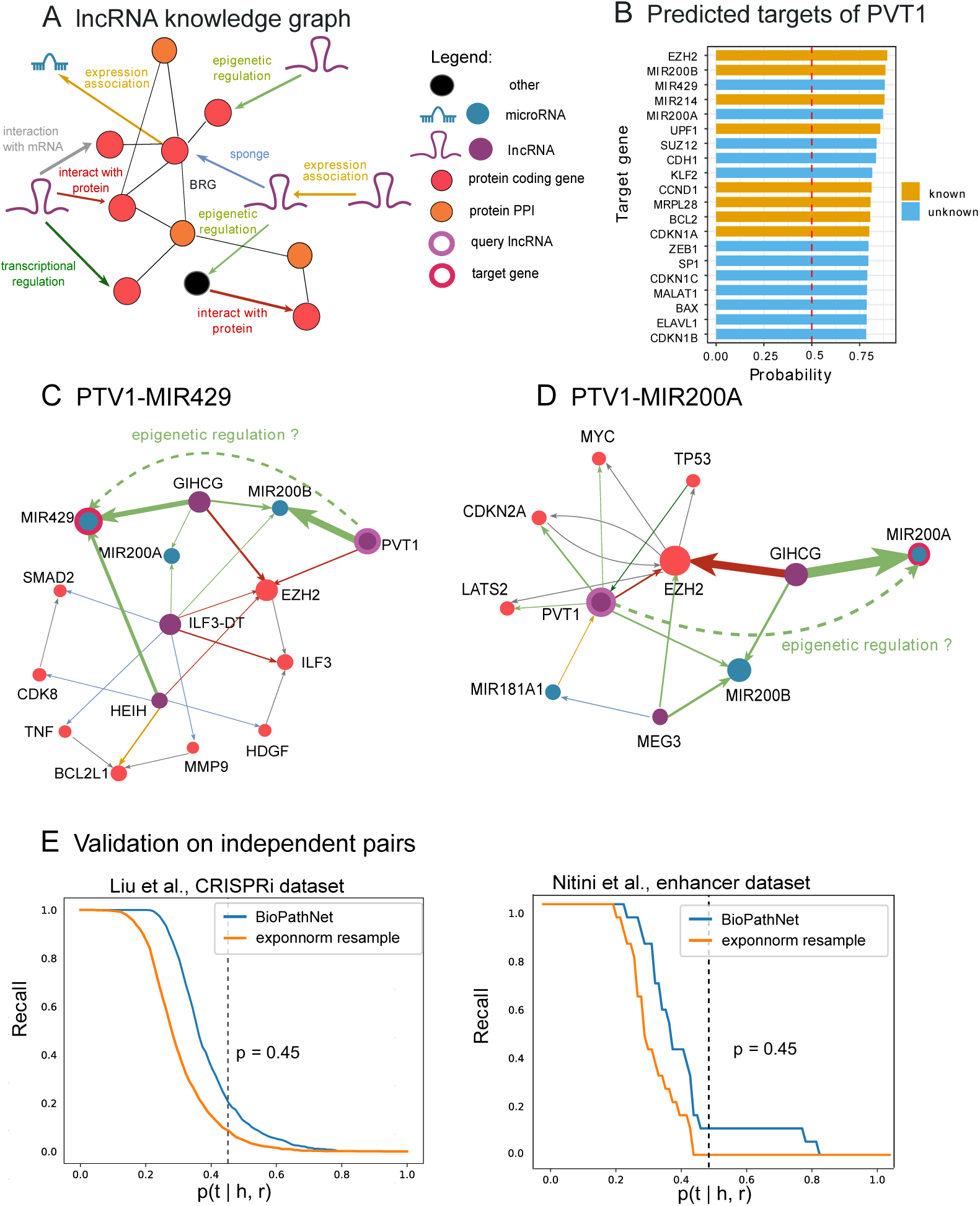
Prediction of novel lncRNA-target regulatory interactions. A) Depiction of the lncRNA-mediated regulation knowledge graph (KG) constructed from the LncTarD 2.0 KG, augmented by incorporating a protein-protein interaction (PPI) network as a BRG for message passing. The graph features six types of node entities: lncRNAs, microRNAs, mRNAs, pseudogenes, transcription factors, proteins, and protein PPI, the latter representing genes from the external PPI network not originally included in LncTarD 2.0. Various types of regulatory relationships are indicated by directed edges of different colors, while protein-protein interactions from the BRG are shown with black undirected edges. B) BioPathNet predicted targets for the cancer lncRNA PVT1, ranked by prediction probability. Annotated targets are depicted in orange, while novel interacting partners are depicted in light blue. C-D) Explanations for the top two predicted PVT1’s novel targets, MIR429 and MIR200A. The top 10 most crucial paths for prediction, ranked by gradient, are shown for both examples. The edge width represents the frequency of appearance in paths; therefore, that connection is important for the prediction. Edge colors indicate the different regulatory mechanisms, following the color code of Figure 6A. E) Independent evaluation of predictions based on external datasets from CRISPR and enhancer-based experiments.

**Table 1:**
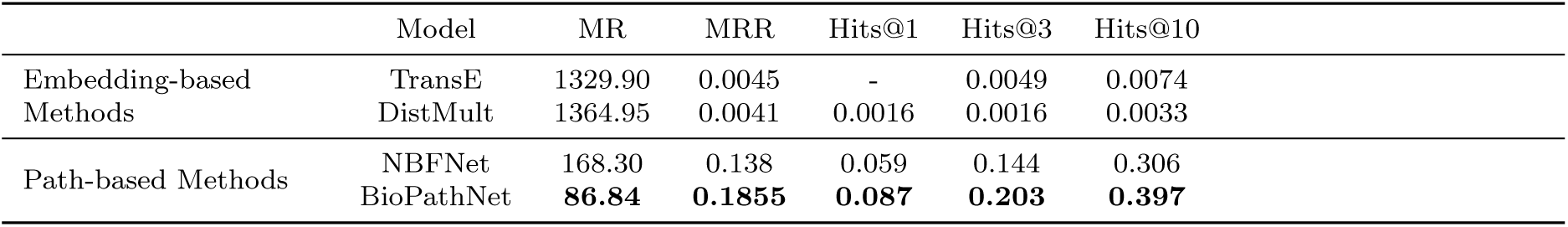
Performance comparison of embedding-based and path-based knowledge graph completion methods in terms of MR, MRR, and Hits@k for k = 1, 3, and 10.

**Table 2:**
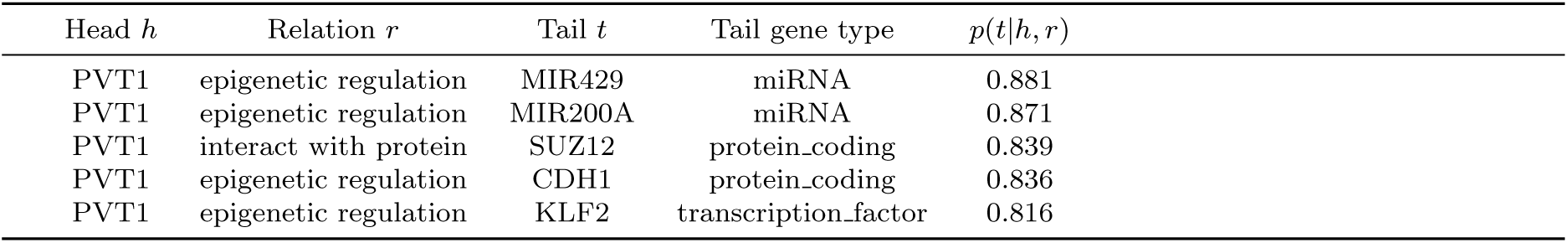
Top 5 novel predicted regulations of PVT1 with highest conditioned probabilities.

In the past year, novel datasets of potential lncRNA-target interactions have been generated, shedding light on the regulatory mechanisms of lncRNAs. In the absence of a gold standard, in order to evaluate the predictive power and generalization capacity of BioPathNet and its underlying KG on the lncRNA-target prediction task, we assessed the method’s recall on two datasets, treating them as independent test sets. The first dataset comprises lncRNA target genes showing significant perturbation in a study by Liu et al. [101], which involved the interference of thousands of lncRNA loci using CRISPRi. In this study, the authors reported target gene-lncRNA regulation pairs. The second smaller dataset encompasses a set of enhancer-like lncRNAs and their potential cis targets determined via chromatin interactions, as defined in Ntini et al. [102]. We evaluated the method by determining how many novel interactions from these datasets (i.e., those not included in the KG) could be identified at different probability thresholds, thereby constructing a recall curve for each dataset. By comparing these results to a random scenario, where the conditional probabilities of the potential novel pairs were randomly sampled from a background distribution (see Methods), we observed that the recall of true lncRNA-target gene pairs in both datasets exceeded that of the recall curve derived from random pairs. This indicates that BioPathNet, trained on the lncRNA-target gene prediction task, can score new datasets containing potential new regulatory interactions significantly better than random.

## 3 Discussion

Biomedical KGs structure information by representing entities (genes, proteins, diseases, drugs) as nodes and their relationships (interactions, associations, regulations) as edges. They integrate diverse data types and enable complex network analysis. Despite high-throughput experiments, many relationships in these graphs remain undiscovered. Link prediction (LP) methods are crucial for inferring missing or potential associations by analyzing network topology.

In this work, we introduce BioPathNet, a message-passing neural network designed to leverage the power of path representation learning for link prediction on biomedical KGs. It is based on the NBFNet algorithm, which efficiently enumerates optimal paths between nodes with the Bellman-Ford algorithm and propagates subpath representations via message passing. BioPathNet introduces several advancements, including the use of a background regulatory graph (BRG) for improved message passing and a node-aware negative sampling strategy to improve learning precision and address graph heterogeneity, design choices that were crucial to improving the performance of specific tasks.

As a proof of concept for biological applications, we evaluated BioPathNet’s ability to reconstruct KEGG-gene annotations for gene function prediction. BioPathNet outperformed node embedding methods, including graph neural networks, achieving over 20% improvement compared to KGE models (TransE, DistMult, RotatE) and over 30% compared to R-GCN without BRG, with a 50% improvement using BRG. This demonstrates BioPathNet’s superior ability to leverage biological regulation information for accurate pathway and gene prediction. We believe BioPathNet outperformed embedding methods by exploiting path-based reasoning to learn representations between nodes based on path relations rather than optimizing one-hop relations. BioPathNet prioritizes relational paths between key entity groups, essential for biological applications and noisy KGs. The BRG in BioPathNet enhanced gene functional annotation by leveraging rich regulatory relationships, enabling comprehensive (gene, pathway) pair representations.

For a more challenging task of predicting drugs for disease treatment, we applied BioPathNet to the zero-shot prediction scenario defined by the state-of-the-art method TxGNN. BioPathNet outperformed TxGNN across all five zero-shot disease splits (adrenal gland, anemia, cardiovascular, cell proliferation, and mental health), with an average AUPRC increase of 20.2%, demonstrating the effectiveness of path-based reasoning in predicting indications. Additionally, BioPathNet achieved higher Recall@k values, prioritizing known treatments better in the top predictions. Specifically, BioPathNet recovered 61.9% of known treatments at k=20, compared to 53.9% with TxGNN. This is especially valuable in biology, as BioPathNet’s enhanced prioritization reduces the number of predictions requiring verification for biological plausibility during hypothesis generation or experimental validation.

In predicting drug contraindications, BioPathNet showed comparable results to TxGNN but with slightly higher performance variance. This variability likely arises because TxGNN relies on stable auxiliary node embeddings for disease similarity, while BioPathNet does not. Instead, BioPathNet makes predictions based on paths connecting disease entities and target drugs so that each disease split might present a different set of edges after removing 95% of connections in zero-shot learning, thus introducing more variability during inference. When comparing BioPathNet and node embedding methods like TxGNN, other advantages and limitations become apparent. TxGNN requires a pre-training phase, using all edges to learn node embeddings equally, followed by fine-tuning with specific relations (indication, contraindications) focusing on drug and disease nodes. It also enhances disease nodes with limited molecular characterization using a gated auxiliary embedding based on node degree. In contrast, BioPathNet uses non-drug-disease triplets for message passing within a BRG but does not require separate pre-training and fine-tuning phases. This simplifies and accelerates training and adds flexibility to our method, allowing the use of different background regulatory graphs without the need for pretraining from scratch. However, the higher variance of BioPathNet compared to TxGNN may be due to the lack of pre-training, as pre-training helps reduce variance by providing a general understanding of relevant features, leading to more stable predictions.

Path embedding methods like BioPathNet enhance representations with multi-hop relationships and offer greater interpretability than node embeddings by tracing and visualizing paths, as well as influent nodes, which aids in verifying predictions and hypothesis generation. Incorporating a Biological Regulatory Graph (BRG) further improves path expressiveness and interpretability, revealing crucial paths and validating predictions. For instance, BioPathNet’s path gradients clarify drug-disease associations, such as Bosutinib for ALL and Acitretin for gastric cancer, and highlight key paths and genes like SMC1A and POLA1 in Clofarabine’s mechanism. Node embedding methods lack intrinsic interpretability and insights into paths or relationships, requiring a post-hoc interpretability framework as seen in TxGNN, yet they are straightforward to comprehend. Node embedding creates high-dimensional vector representations that are applicable in downstream tasks, with nodes closer in embedding space, reflecting their similarities via methods like t-SNE or UMAP. In contrast, path embedding, despite capturing a richer context, is more abstract and less straightforward for downstream applications.

While evaluating BioPathNet against TxGNN in zero-shot scenarios for disease-drug predictions, we observed that TxGNN’s data splits resembled near zero-shot scenarios. Some connections between drugs and diseases similar to the target disease were retained in the training graph, possibly leading to information leakage during inference. Despite both methods being evaluated on the same data splits, we wanted to determine if BioPathNet could still predict meaningful disease-drug indications when these informative edges were intentionally excluded from the inference graph. As an example, consider the case of Clofarabine, a known indication for ALL, also annotated in the PrimeKG database (Supplementary Figure 2). If the connection between ‘leukemia, lymphocytic, susceptibility to’ and Clofarabine is not removed during training and inference, the model can reconstruct the link between leukemia (disease) and Clofarabine through this path, exploiting the similarity between ‘leukemia, lymphocytic, susceptibility to’ and ‘leukemia (disease)’. To improve the interpretability, we removed the link during inference: the model this time reveals important nodes such as genes SMC1A, involved in chromosome cohesion during cell division and DNA repair, and POLA1, part of the DNA polymerase alpha subunit. Given that Clofarabine is a purine nucleoside metabolized intracellularly to inhibit DNA synthesis [67, 103], the model identifies key components of the drug’s mode of action through alternative paths. A similar example is shown for gastric cancer (Supplementary Figure 2): to reconstruct the link between gastric cancer and its known indication Capecitabine, BioPathNet initially uses the path containing the retained connection between a similar disease, ‘gastric linitis plastica,’ and Capecitabine. When we remove this link during inference, BioPathNet cannot rely on disease similarities and must find another path to obtain the same prediction. In summary, using a custom or modified graph during inference, with the removal of diseases similar to the disease of interest, highlights the flexibility of path-based methods in adapting to graph structure changes, unlike embedding-based approaches reliant on direct node connections. This experiment suggests the potential of using different inference graphs and towards inductive reasoning settings, particularly beneficial in scenarios with new nodes emerging during inference. Future research will delve deeper into fully inductive reasoning tasks.

We demonstrate the versatility of BioPathNet in addressing diverse problems across various KGs. For instance, we show that BioPathNet can be confidently used in path-based reasoning and explainable predictions of SL gene pairs and can identify novel SL pairs crucial for improving cancer treatment efficacy. By leveraging heterogeneous graph information and node-type-specific negative sampling, BioPathNet achieves precise SL predictions, often surpassing state-of-the-art methods like KR4SL.

The task of inferring novel lncRNA-mRNA regulatory relationship is the hardest in this context, as few and noisy data are available for training, and the KG is very sparse compared to other settings. Here we attempt this for the first time this task making use of a lncRNA-gene-specific KG coupled with a background regulatory graph (BRG) for enhanced message passing, similar to the other tasks. Despite an MRR of 0.19 - lower than tasks like drug repurposing or synthetic lethality — indicating that BioPathNet could probably benefit from more training data, BioPathNet still strongly learns the structure of the lncRNA-mRNAs regulatory graph, and it is much more effective than node-embedding methods in reconstructing true lncRNA-mRNA relationships, by leveraging multiple paths in the sparse graph, and by that compensating for the lack of direct connections.

As a final remark, under the open-world assumption, evaluating model performance on incomplete KGs may not fully reflect their capabilities. Metrics, like MRR, can degrade in scenarios with high incompleteness, where missing links correlate with specific entities, as discussed by Yang et al. [104]. Biomedical KGs exhibit uneven gaps in knowledge distribution, influenced by factors like prevalence and complexity, potentially underestimating BioPathNet’s performance, particularly in tasks such as lncRNA-mRNA prediction.

Limitations of BioPathNet include potential biases in the training data. For example, while BioPathNet successfully retrieved almost all known treatments for Alzheimer’s disease (AD) in its top predictions, it missed FDA-approved drugs such as Pramiracetam, Memantine, and Lecanemab, which were not listed in the PrimeKG database and lacked disease indications. As a result, the model couldn’t learn their connections and did not identify them as potential treatments. Predictions for known symptom-treating drugs focused on neuropsychiatric-related diseases, but emphasizing molecular interactions could uncover more disease-modifying treatments. Future improvements might involve excluding message passing over dominant relations like indications and prioritizing molecular interactions to elucidate mechanisms underlying less understood diseases like Alzheimer’s.

## 4 Conclusion

In conclusion, BioPathNet is a novel method for link prediction on biological KGs using path embedding. It excels in gene function prediction, zero-shot drug indication, synthetic lethality pair, and mRNA-lncRNA interaction tasks, consistently outperforming state-of-the-art methods. Its interpretability framework retrieves and visualizes key prediction paths, enhancing understanding, uncovering biases, and evaluating biological plausibility. Future work could focus on evaluating BioPathNet in inductive settings, refining the KG with more informative sources, and fine-grained relations. Utilizing condition-specific KGs enriched with detailed tissue, patient, pathway, and disease knowledge from platforms like BioCypher [105] could enhance reasoning capabilities. Additionally, integrating node features in path representations, such as experimental sequencing data, could further improve predictions. We believe that in the future BioPathNet could pave the way for foundational models in link prediction within biomedical KGs, significantly advancing the pace of hypothesis generation across various biological and biomedical domains.

## 5 Methods

### 5.1 Knowledge Graph Completion

A knowledge graph *KG* = {(*u, r, v*)}*_u,v ∈ E,r ∈ R_* is a heterogeneous directed graph with entities *E* as nodes, relations *R*, and a list of triplets (*u, r, v*) that represent the edges in the graph. Here, *u* (head) and *v* (tail) are entities, and *r* (relation) is an edge or link. The graph is considered heterogeneous because different entities may have different types, e.g. a node representing a gene versus a node representing a disease. The graph is directed because (*u, r, v*) being in the KG does not imply (*v, r, u*) is contained as well. Knowledge graph completion involves predicting missing the missing links, i.e. triplets, categorized into three tasks: 1) Tail prediction (*u, r,* ?) - predicting the tail entity given the head entity and the relationship; 2) Head prediction (?*, r, v*) - predicting the head entity given the relationship and the tail entity; 3) Relation prediction (*u,* ?*, v*) - predicting the relationship given the head and tail entities [106].

### 5.2 Neural Bellman-Ford Network (NBFNet)

Our newly developed BioPathNet is a path-representation learning-based method for graph completion built upon the NBFNet framework [50]. Unlike node embedding methods or node GNN encoders that infer links between entities in a KG by learning node representations in an embedding space, NBFNet is a general graph neural network framework that performs link prediction by learning representations for each path from the query entity *u* to potential tail entities *v*. More specifically, in NBFNet, the path formulation is represented by a generalized sum of path representations between *u* and *v* (see Extended Methods, Supplementary File).

Two key factors contribute to NBFNet’s scalability for large graphs and its effectiveness in learning tasks: the use of the generalized Bellman-Ford dynamic programming framework for path representation and the abstraction of this process into a neural formulation.

### Generalized Bellmann-Ford path representation

To achieve a scalable path formulation, NBFNet utilizes a generalized version of the Bellman-Ford dynamic programming algorithm [107]. This generalization transforms the original Bellman-Ford algorithm for shortest path calculation into a versatile framework that simultaneously computes pair representations *h_q_*(*u, v*) for a given entity *u*, query relation *q*, and all vertices *v* in a graph. This approach reduces the computational cost to polynomial time relative to the number of nodes and edges in the graph.

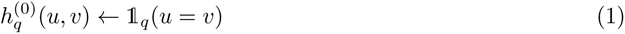

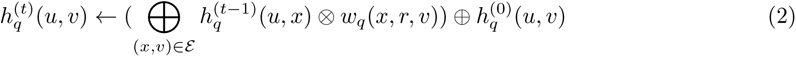

In this formulation, the first equation initializes the boundary condition on the source node (equation 1), representing the shortest path between *u* and *v* at the start. If the head and tail nodes coincide (*u* = *v*), the boundary condition is set to the generalized 1, which corresponds to 0 in the shortest path context (i.e., the shortest distance between a node and itself is zero) and to ∞ in the case *u* ≠ *v*. Equation 2 describes the Bellman-Ford iteration, updating the shortest path distance between *u* and *v*. In each iteration, the representation from the previous layer (*t* − 1) is multiplied by the transition edge representation *w_q_* to obtain the new representation *h_q_*(*u, v*). The algorithm propagates the boundary condition from the source node to its neighbors. It is important to note that there is a distinction between query relation *q* and the relation *r* in the graph. The query relation *q* is used to initialize the source node (boundary condition), while then the transition edge representations *w_q_*(*x, r, v*) are obtained by the multiplication of relation *r* in the graph. Both embeddings are learned. Using the distributive properties of multiplication, all prefixes are computed simultaneously. This iterative process continues, assessing potential target nodes, until all paths from the source to the tail node are covered after *t* iterations, where *t* is the path length. For a more detailed description, refer to the Extended Methods in the Supplementary File.

### Neural formulation

By abstracting the boundary condition in equation 1 to an indicator function, the multiplication operator in equation 2 to a message passing formulation, and the summation operator to a general aggregation function, NBFNet extends the generalized path formulation of the Bellman-Ford algorithm into a graph neural network framework.

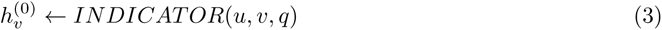

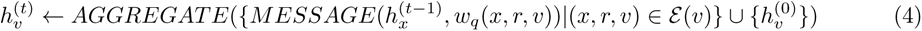

For the indicator function, NBFNet learns the query relation embedding *q* and assigns *q* to node *v* if *v* equals the source node *u*. For message passing, it uses relational operators from KG embeddings: TransE (translation), DistMult (multiplication), and RotatE (rotation). Aggregation functions are permutation-invariant functions from GNN literature, including sum, mean, max, and principal neighborhood aggregation (PNA). Instead of traditional edge representations like transition probabilities or lengths, NBFNet parameterizes edge representations as a linear function of the query relation [50].

NBFNet can be interpreted as a novel GNN framework for learning pair representations. Unlike typical GNNs, which compute pair representations as independent node embeddings *h*(*u*) and *h*(*v*), NBFNet conditions each node’s representation *h_q_*(*u*) and *h_q_*(*v*) on the source node and query relation *q*. The resulting pair representation *h_q_*(*u, v*) is then used for link prediction, predicting the tail entity *v* given the head entity *u* and relation *q*. This is formulated as the conditional likelihood of the tail entity *v* as:

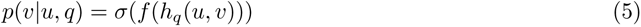

where *σ*(_) is the sigmoid function and *f* (_) is a feed-forward neural network.

### 5.3 BioPathNet - Biomedical Knowledge Graph Completion

Considering the unique characteristics of biological KGs, we introduce BioPathNet, a graph neural network framework based on NBFNet [50]. BioPathNet is designed to predict missing links in biomedical KGs and is applied to four biological tasks:

1. *Gene Function Prediction*: Identifying potential novel functions for genes via gene-pathway associations.
2. *Drug Repurposing* : Discovering new indications for existing drugs by analyzing drug-disease associations for established drugs.
3. *Synthetic Lethality Prediction*: Identifying novel synthetic lethality gene pairs.
4. *lncRNA-Gene Target Prediction*: prediction of regulatory relationships between lncRNAs and their putative target genes.

#### Path representation in BioPathNet

As path-based reasoning method, path representations *h_q_*(*u, v*) in BioPathNet are learned starting at a source node *u* to all potential target nodes *v* based on the relations *r* along the path, following the NBFNet parametrization (equations 3 and 4) but with important enhancements which make BioPathNet more suited for biological KGs. Firstly, we make use of entity type information, which is not used in NBFNet originally. Secondly, we pool additional data sources beyond the target KG to augment the knowledge available during reasoning for the target link prediction task. Specifically, given a KG *G*_1_ for which we wish to predict the missing links, we add an additional graph *G*_2_, as a Biological Regulatory Graph (BRG), into the path representation computation. These augmentation to the original NBFNet method constitute our BioPathNet.

The incorporation of an external BRG, e.g. protein-protein interaction or gene regulatory network, provides additional edges (knowledge) that are used solely for message passing, enhancing the prediction of links of interest (Figure 1C). For example, in predicting the missing link (*u, r, v*), where (*u, r, v*) always comes from the target KG *G*_1_, messages can be passed only along paths in *G*_1_ yielding a prediction path such as in Figure 1D. Alternatively, a BRG *G*_2_ can be supplied to further information about the predictions, leveraging other knowledge bases (e.g., including relations between type 2 and 3 nodes), as illustrated in Figure 1E.

Consequently, in BioPathNet, equation 4 is modified to always take (*u, r,* ?) from *G*_1_ but do aggregate and send messages across all edges *G*_1_ ∪ *G*_2_. In equation 4 the edges (*x, r, z*) ∈ E(*v*) come from *G*_1_ ∪ *G*_2_ rather than just *G*_1_.

After performing link prediction with the modified message passing scheme, BioPathNet ranks all candidate tail entities according to their likelihood *p*(*v*|*u, q*) to form a true triplet with a given head entity and relation *q* as the query. During the training of the model in a supervised setting, the negative log-likelihood is minimized between positive samples ⟨*u, r, v*⟩ (i.e., known triplets composed of a head node and tail node and the relationship between them) and negative samples ⟨*u, r, v^′^*⟩, which are generated by corrupting *v* (i.e., substituting the true *v* with another node *v*’).

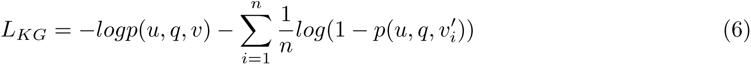

where *n* is the number of negative samples per positive sample and (*u, q, v^′^*) is the *i*-th negative sample for KGs. The same approach is used for the prediction of *v* given *u* and *r^−^*^1^ for the reverse relation of *r*. Unlike the original NBFNet, BioPathNet implements an entity-type aware negative sampling scheme.

This means that when sampling negative *v*’ for training, we consider the node type and sample *v*’ only from the same type as *v*. This approach dramatically reduces the sample space and allows the model to learn better decision boundaries by focusing on sufficiently difficult negative samples. Importantly, the additional edges from the BRG are not used for sampling positive and negative triplets, ensuring that the computation of the loss in equation 6 remains unchanged.

### 5.4 Interpretation of prediction - Visualization of most important paths

Leveraging the NBFNet framework, BioPathNet predictions can be directly interpreted through paths. This feature is crucial for biomedical tasks, where understanding the mechanisms behind each prediction is essential. These interpretations highlight the paths that most significantly contribute to the prediction *p*(*v*|*u, q*). Using local interpretation methods, we approximate the local landscape of BioPathNet with a linear model over the set of all paths, and the importance is then defined by its weight in the linear model, which can be computed as the partial derivative of the prediction with respect to the path [50]. Formally, the top-k path interpretations for *p*(*u, q, v*) are defined as:

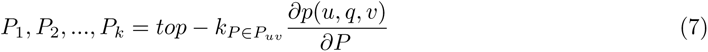

While directly computing the importance of all paths is intractable, NBFNet approximates them with edge importance. Specifically, the importance of each path is approximated by the sum of the importance of edges in that path, and therefore intuitively, the top-k path interpretations are equivalent to the top-k longest paths on the edge importance graph.

For the visualization plot, we consider the top 10 most important paths ranked by gradient, with the edge width reflecting the number of times an edge appears in paths. Furthermore, the most important path is highlighted in red. In summary, our interpretability allows predictions to be assessed by their biological plausibility for hypothesis generation or validation in the laboratory.

### 5.5 KG construction, Data pre-processing, and BioPathNet training

#### Gene function prediction task

For this task, we used the knowledge graph (KG) from the KEGG database (G1), extracted from ConsensuPathDB, to train the model on gene (G) - pathway (P) interactions. Additionally, we utilized a BRG containing regulatory relationships between gene-gene (G-G), gene-chemical (G-C), and chemical-chemical (C-C) obtained from Pathway Commons (G2)[52, 53]. These interactions were represented as triplets in the format (node1, relationType, node2), such as (BABAM1, interacts-with, PSMD14). Details of the data, graph, relation types, and train, validation, and test sets are provided in Supplementary Tables 1 and 2. During data pre-processing, we removed KEGG pathways with fewer than 10 annotations to genes. Next, we loaded both the BRG and KEGG graph (of pathways and genes [P-G]) as a multi-graph in the network, maintaining only the biggest connected component, thereby removing 11 nodes present in components of only 2–3 nodes. We trained BioPathNet using 70% of the P-G triplets, which were randomly split, with and without incorporating the underlying BRG as a message-passing graph. We used 10% of the P-G triplets for validation, and the remaining 20% were reserved for testing. When the BRG was not utilized, we took an additional step to exclude triplets containing genes that appeared in the validation or test sets but were absent from the training set. Hyperparameters were optimized based on the validation MRR, resulting in an optimal set of parameters for downstream analysis (Supplementary Table 3).

#### Drug repurposing task

For this task, we used the PrimeKG database, an extensive multi-modal knowledge graph designed to integrate and unify diverse types of resources and biomedical and clinical data, such as gene-gene interactions, gene-disease associations, and drug-disease information [60] (Supplementary Figure 1). A summary of the node and edge relations can be found in [59], and details of the number of graph’s nodes and edges used for message passing, training, validation, and test are provided in Supplementary Table 4.

For training and evaluating BioPathNet we used the same data and data splits, as defined by TxGNN, a geometric deep learning model for zero-shot drug repurposing predictions also based on PrimeKG. Five distinct zero-shot disease areas were used: adrenal gland disorders, anemia, cardiovascular diseases, cell proliferation issues, and mental health conditions (Supplementary Table 4). A disease area encompasses a specific group of related diseases. For instance, the “cell proliferation” area includes various cancer types. We utilized TxGNN’s data split to create training, validation, and testing datasets reflecting a zero-shot prediction scenario in a proportion of 0.83:0.12:0.05. By using the TxGNN code (https://github.com/mims-harvard/TxGNN from Apr 13, 2023, commit “1000aac”), the different splits were created by removing all triplets with the relation types indication and contraindications for a disease area from the training dataset, along with 95% of connections to biomedical entities such as proteins and phenotypes [59]. This split simulates minimal molecular characterization of a disease area combined with no knowledge of therapeutic opportunities. While TxGNN constructs reverse edges, we removed those beforehand, since BioPathNet inherently adds reverse triplets (and the reverse relations) during reasoning.

Only edges between drugs and diseases, such as indication and contraindication, were used to train BioPathNet in a supervised manner (G1 graph). The remainder of the PrimeKG graph served as the BRG for message passing (G2 graph). After removing reverse relations, the BioPathNet model used 5.7 million directed edges for message passing per prediction setting (Supplementary Table 4). These edges, unlike supervised training triplets, were protein-protein or disease-disease relations. The training and validation sets averaged 33,000 and 4,000 edges, respectively, across five disease areas (Supplementary Table 4).

We further created our custom data split with TxGNN’s “disease eval” code to evaluate the performance in predicting drugs for the neurodegenerative disorder Alzheimer’s disease (AD). Drugs that were associated with various AD diseases were moved to the test set (Supplementary Table 5). All models were trained for 10 epochs, employing an early stopping mechanism that retained the best model based on validation set performance (MRR, see below). The final hyperparameters for all five disease area splits are reported in Supplementary Table 6. All experiments were repeated for five different datasplit seeds, using the exact seeds employed by TxGNN to ensure a fair comparison. Each data split seed resulted in slightly different training and validation sets for each disease area due to the random removal of edges to simulate the zero-shot scenario for the disease under study every time. Performance metrics were reported on the test set as the mean ± standard deviation across the five seeds (Supplementary Table 7).

#### Synthetic lethality (SL) prediction task

For training and inference of BioPathNet, we used the SynLethDB-v2.0 [93] data as a KG, an updated database compiling SL relationships derived from screening experiments, as well as computational predictions, providing a comprehensive resource for exploring gene interactions in cancer. BioPathNet was trained on preprocessed data, i.e. the SL gene pairs extracted from SynLethDB-v2.0, as provided by KR4SL, a KG-based model designed to predict SL interactions in cancer [49], and downloaded from their GitHub repository (https://github.com/JieZheng-ShanghaiTech/KR4SL from Dec 8, 2023 - commit “61b5c84”). In detail, the SL gene pairs from SynLethDB-v2.0 for humans were randomly split into train, validation, and test triples in a ratio of 7:1:2, following the data split of KR4SL (Supplementary Tables 8 and 9). The pre-processed data was modified to fit the BioPathNet format by removing the reverse edges introduced by KR4SL for SL gene pairs (following the same pre-processing scheme in the drug repurposing task), as reverse edges are implicitly added by BioPathNet by default.

On top of the SL pairs, SynLethDB-v2.0 constructs a KG including relations between gene entities, pathways, and three types of Gene Ontology (GO) terms: biological processes (BP), molecular functions (MF), and cellular components (CC) augmented using OntoProtein [44]. While SL pairs were used for supervised training (G1), the rest of the KG (G2) was used as BRG for message passing only, following the same scheme from previous tasks (Supplementary Tables 8–10). Hyperparameters were optimized based on the validation MRR, resulting in an optimal set of parameters for downstream analysis (Supplementary Table 11). We trained and evaluated BioPathNet at different confidence thresholds on the SL pairs, ranging from 0.1 to 0.8, as SL pairs in the database have varying confidence levels. This “thresholded data” approach contrasts with “unthresholded data,” which includes all SL pairs from SynLethDB-v2.0 without filtering by confidence score. For a fair comparison, we ran BioPathNet for the same number of epochs using identical seeds and tuned hyperparameters on unthresholded data, which were then applied to thresholded data.

Since SL relationships are symmetric between genes, the final score for a gene *v* to be an SL partner of gene *u* is computed by considering both the feed-forward neural network transformed representation of tail node *v* given head node *u* and the SL relation, *f* (*h_q_*(*u, v*)), as well as the transformed symmetric representation, 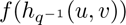 Thus, the final SL score for a gene *v* to be an SL partner of gene *u* from BioPathNet is reported as:

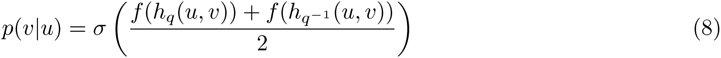

#### LncRNA-gene target prediction task

For this task, we used the LncTarD 2.0 database [108], a manually curated database of 8360 experimentally supported functional lncRNA-target regulatory interactions in human diseases, categorized into seven mechanisms of lncRNA-target regulation: *ceRNA or sponge*, *chromatin looping*, *epigenetic regulation*, *expression association*, *interact with mRNA*, *interact with protein* and *transcriptional regulation*. First, incomplete gene information, such as missing Ensembl IDs or gene names, was resolved via Gencode and HGNC mapping. Second, as only 12 pairs of regulations belonged to the chromatin looping category, we re-labeled them as transcriptional regulation after manually inspecting every regulatory interaction in the scientific literature. This condensed the interaction relationships into six distinct types. A KG was constructed from LncTarD 2.0 (G1), where entities are the genes involved, and relations are the regulatory mechanisms. For a triplet (*u, r, v*), the head *u* corresponds to the regulator (e.g. lncRNA), the relation *r* to the regulation mechanisms, and the tail *v* to the target gene. On top of the LncTarD 2.0 KG, we added the BRG derived from PathwayCommons (G2), the same used for the gene function prediction task. As there is no direct link between small molecules and lncRNA, we only used PPI, discarding other types of relations. This enriches the original KG with additional connectivity. In the end, BioPathNet was trained on the lncRNA interactions from LncTarD 2.0 KG. The specific numbers of nodes and edges for the LncTarD-derived KG, the number of edges and nodes corresponding to the different relation types, as well as those used for train, validation, and testing of BioPathNet are detailed in Supplementary Tables 14 and 15. To enable node type-aware negative sampling, node entities were labeled with six different categories: *lncRNA*, *mRNA*, *microRNA*, *transcription factor*, *protein* and *protein ppi* (this last one to specifically identify nodes from ppi interactions from the BRG, whose edges are only used for message passing and not supervised training). The optimal parameters of BioPathNet in this setting, determined through the MRR on the validation set, are reported in Supplementary Table 16.

Further, we ranked the interaction partners of 42 lncRNAs, including PVT1, sourced from the studies of [102] and [101]. We considered a regulatory relationship between the lncRNA of interest and its target genes when there existed any relation *r* for which the conditional probability exceeded a threshold *t*, *p*(*v*|*u, q*) ≥ *t*. Conversely, if no such relation exists, it is considered that there is no regulatory relationship. The threshold we used here is the average probability of the triples that overlap with the training set outputted by BioPathNet. These probabilities approximated an exponential, normal distribution and were also used for random sampling, constituting a random baseline.

### 5.6 Comparison with baselines

We compared BioPathNet against several baselines. Among the KG Embedding methods, we benchmarked TransE, DistMult and RotatE, belonging to shallow models learning embeddings with an encoder for each relation and node. The latent embedding space is restricted by the semantic relationship *r* between *u* and *v* nodes. RotatE models relations as rotations in the complex plane to capture symmetric and antisymmetric patterns [37], TransE represents relations as translations between entities [71], and DistMult uses diagonal matrices to capture symmetric relationships through element-wise multiplication of entity and relation embeddings [38]. We also benchmarked BioPathNet against the Graph neural network-based R-GCN, a method that performs both node classification and link prediction tasks, extending traditional GCNs to handle multi-relational data by introducing relation-specific weight matrices [109]. It updates node representations by aggregating information from neighbors, considering the type of edge connecting them, which allows it to capture the distinct characteristics of different semantics of each relation within a graph.

For the Drug repurposing prediction task, we compared BioPathNet to TxGNN, a state-of-the-art model for predicting drug-disease relationships in zero-shot scenarios, where minimal prior information or treatment history is available [60]. Leveraging PrimeKG [60], a comprehensive biomedical knowledge graph, TxGNN uses R-GCNs to learn embeddings of drugs and diseases, capturing complex interactions by mapping them into a shared latent space.

For the Synthetic lethality pair prediction task, BioPathNet was benchmarked against Knowledge Representation for Synthetic Lethality (KR4SL), a path-representation learning GNN-based method designed specifically for the explainable prediction of SL gene pairs in cancer [49].

All baseline methods were re-trained with optimal parameters to ensure a fair comparison. Detailed descriptions of each baseline and the specifics of the final model parameters are provided in the Extended Methods section of the Supplementary File.

### 5.7 Model evaluation

Various metrics were used to evaluate BioPathNet across different tasks and compare its performance with baseline methods. For all tasks, methods were evaluated based on: Mean Rank (MR), the average rank of the true positive among all predicted candidates; Mean Reciprocal Rank (MRR), the average of the reciprocal ranks of the first relevant item. Hits@*k*, the proportion of true positives ranked within the top *k* predictions. Values for these metrics range in [0, 1], and the larger the value, the better the model (for an extensive explanation of these metrics, refer to the Extended Methods section of the Supplementary File).

While KGC models the conditional probability of predicting the tail entity *v* given the head entity *u* and relation *r*, evaluating the joint probability of *u*, *v*, and *r* may be more comprehensive. To ensure consistency with TxGNN in drug prediction, we also used AUPRC to summarize precision and recall across thresholds, along with specificity and F1 score at a 0.5 threshold, using TxGNN’s evaluation code. This approach assesses the performance of each disease node. For details on computing AUPRC in the comparison between BioPathNet and TxGNN, refer to the Extended Methods section in the Supplementary File.

For the SL prediction task, we compared the seed-wise performance of our model with the performance of KR4SL using metrics inherent to the KR4SL framework’s code, specifically NDCG@k, Recall@k, and Precision@k (see Extended Methods, Supplementary File). Moreover, we computed MRR for both BioPathNet and KR4SL by first calculating MRR for each query gene and then averaging gene-wise MRRs overall query genes.

## 6 Data Availability

Data for the gene function prediction task can be downloaded from the public platforms of PathwayCommons and ConsensusDB. We refer to the methods for more instructions. In the drug-disease prediction task, PrimeKG’s data can be automatically downloaded and data splits generated by TxGNN. SynLethDB data was processed by KR4SL, which we obtained through their GitHub repository. Data for lncRNA target prediction was obtained over LncTarD 2.0. Further preprocessing was done to fit BioPathNet’s data format, all scripts can be found in https://github.com/emyyue/BioPathNet.

## 7 Code Availability

The BioPathNet model and all the code necessary for reproducing our results is publicly available via GitHub at https://github.com/emyyue/BioPathNet. An archived version will be deposited in the Zenodo database upon acceptance.

## Acknowledgments

We thank the Helmholtz Association under CausalCellDynamics (to A.M., Y.H., S.F. and S.X.) and the joint research school ‘Munich School for Data Science (MUDS)’ (to S.O. and S.F.); the Joachim Herz Foundation (to Y.H.); the CNATM BMBF to A.M. M.A. received funding from the National Institutes of Health/National Institute on Aging through grants RF1AG059093, U01AG061359, and R01AG081322.

## Author contributions

Y.H., S.X., Z.Z., A.M., and J.T. conceived the project. Y.H. implemented the BioPathNet model with help from S.X.and Z.Z. and carried out the experiments on gene function prediction and drug repurposing, with help from S.F.. S.O. performed the full KR4SL analysis and comparison with BioPathNet, Y.H. and S.F. performed the TxGNN analysis and comparsion with BioPathNet; H.C. perfomed the lncRNA-target prediction analysis with help from Y.H.. M.U., M.C.T, and M.A. helped with the interpretation of the results. A.M. and S.X. co-supervised the study; Y.H., S.O., and A.M. wrote the manuscript with help from S.F. and S.X. and additional input from all co-authors. All authors reviewed and approved the final manuscript.

