## Supplementary file for "Path-based reasoning for biomedical knowledge graphs with BioPathNet"

[+]equal contribution.
[§]co-last author.
[1]Computational Health Center, Helmholtz Center Munich, Ingolstaedter Landstrasse 1, Neuherberg, 85764, Bavaria, Germany.
[2]School of Life Sciences, Technical University of Munich, Alte Akademie 8, Freising, 85354, Bavaria, Germany.
[3]Department, Mila - Québec AI Institute, 6666 St-Urbain, Montréal, QC H2S 3H1, Quebec, Canada.
[4]Department, Université de Montréal, 2900, boul. Édouard-Montpetit, Montréal, QC H3T 1J4, Quebec, Canada.
[5]School of Computation, Information and Technology, Technical University of Munich, Arcisstrasse 21, Munich, 80333, Bavaria, Germany.
[6]Faculty of Biology, Ludwig-Maximilians University, Grosshaderner Str. 2, Planegg-Martinsried, 82152, Bavaria, Germany.
[7]Department, CIFAR AI Chair, 661 University Ave, Toronto, ON M5G 1M1, Ontario, Canada.
[8]Department, HEC Montréal, 3000 Chem. de la Côte-Sainte-Catherine, Montréal, QC H3T 2A7, Quebec, Canada.

Contributing authors:;

**Keywords:** biomedical knowledge graph, link prediction, graph neural network

### 1 Extended Methods

#### 1.1 Neural Bellman-Ford Network (NBFNet)

Our newly developed BioPathNet is a path-representation learning-based method for graph completion built upon the NBFNet framework. Below, we describe the foundations of NBFNet to provide an understanding of how BioPathNet operates.

Unlike node embedding methods or node GNN encoders that infer links between entities in a KG by learning node representations in an embedding space, NBFNet is a general graph neural network framework that performs link prediction by learning representations for each path from the query entity to potential tail entities. This is done by considering the relations along each possible path between the query entity and the potential tail entity. Given a KG, NBFNet learns to predict the tail node for a query $(u, q, ?)$. It does this by learning a node pair representation $h_q(u, v)$ for nodes $u$ and $v$, which captures all the possible paths between $u$ and $v$ conditioned on $q$. In NBFNet, the path formulation is represented by a generalized sum of path representations between u and v, denoted by $h_q(P)$, with a commutative summation operator $\bigoplus$:

$$h_q(u, v) = h_q(P_1) \oplus h_q(P_2) \oplus ...... \oplus h_q(P_M) \tag{1}$$

where $M$ denotes the possible number of paths between $u$ and $v$. Each path representation $h_q(P)$ is defined as the generalized product of the edges (or edges representations) belonging to that path, with the multiplication operator $\bigotimes$.

$$h_q(P) = w_q(e_1) \otimes w_q(e_1) \otimes ....... \otimes w_q(e_p) \tag{2}$$

where $p$ is number of edges $e_1, ..., e_p$ belonging to path $P$. In compact form, the path formulation can be written as:

$$h_q(u, v) = \bigoplus_{p \in P_{u,v}} \bigotimes_{e \in P} w_q(e) \tag{3}$$

This, in practice, means that we compute the representation between nodes $u$ and $v$ under query relation $q$ by considering every path between the source node $u$ and the target node $v$. Each path is represented by the product of edge representations $w_q$ along that path, and we sum these products to obtain the final representation.

Two key factors contribute to NBFNet's scalability for large graphs and its effectiveness in learning tasks: the use of the generalized Bellman-Ford dynamic programming framework for path representation and the abstraction of this process into a neural formulation.

**Generalized Bellmann-Ford path representation** The approach above, which predicts a link between a head and tail node by enumerating all possible paths and summing their contributions, is not scalable. As the network grows, the number of possible paths between the head and tail nodes increases exponentially, making this method impractical for larger networks. To achieve a scalable path formulation, NBFNet utilizes the Bellman-Ford dynamic programming algorithm, which efficiently finds the shortest path from a single source node to all other nodes in a weighted graph through a recursive process [1]:

$$d[v] = min(d[u] + w(u, v)) \tag{4}$$

By extending this equation to generalize the addition operator to any summation operator $\bigoplus$ and the minimum operator to any multiplication operator $\bigotimes$, we derive the generalized Bellman-Ford algorithm [2]. This generalization transforms the original Bellman-Ford algorithm for shortest path calculation into a versatile framework that simultaneously computes pair representations $h_q(u, v)$ for a given entity $u$, query relation $q$, and all vertices $v$ in a graph. This approach reduces the computational cost to polynomial time relative to the number of nodes and edges in the graph. The formulation of the generalized Bellman-Ford algorithm is as follows:

$$h_q^{(0)}(u,v) \leftarrow \mathbb{1}_q(u=v) \tag{5}$$

$$h_q^{(t)}(u,v) \leftarrow \Big(\bigoplus_{(x,v)\in\mathcal{E}} h_q^{(t-1)}(u,x) \otimes w_q(x,r,v)\Big) \oplus h_q^{(0)}(u,v) \tag{6}$$

It first initializes the boundary condition on the source node (equation 5), representing the shortest path between $u$ and $v$ at the start. If the head and tail nodes coincide ($u = v$), the boundary condition is set to the generalized $\mathbb{1}$, which corresponds to 0 in the shortest path context (i.e., the shortest distance between a node and itself is zero) and to $\infty$ in the case $u \neq v$. Equation 6 describes the Bellman-Ford iteration, updating the shortest path distance between $u$ and $v$. In each iteration, the representation from the previous layer $(t-1)$ is multiplied by the transition edge representation $w_q$ to obtain the new representation $h_q(u,v)$. Here, $W_q$ is a function of the relation between $u$ and $v$. The boundary condition is added at each step to ensure accurate path formulation. The algorithm starts by propagating the boundary condition from the source node to its neighbors. Thanks to the distributive properties of the multiplication operator, all prefixes sharing this propagation are computed simultaneously. This iterative process continues, assessing potential target nodes, until all paths from the source node to the tail node are covered after $t$ iterations, where $t$ represents the path length.

**Neural formulation** The generalized summation and multiplication operators are handcrafted. By abstracting the boundary condition in equation 5 to an indicator function, the multiplication operator in equation 6 to a message passing formulation, and the summation operator to a general aggregation function, NBFNet extends the generalized path formulation of the Bellman-Ford algorithm into a neural network framework. In this formulation, neural network functions parametrize the boundary condition, multiplication, and summation functions, allowing a generalized GNN framework to learn these three neural functions effectively.

$$h_v^{(0)} \leftarrow INDICATOR(u,v,q) \tag{7}$$

$$h_v^{(t)} \leftarrow AGGREGATE(\{MESSAGE(h_x^{(t-1)}, w_q(x,r,v)) | (x,r,v) \in \mathcal{E}(v)\} \cup \{h_v^{(0)}\}) \tag{8}$$

For the indicator function, NBFNet learns the embedding of the query relation $q$ and assigns $q$ to node $v$ if $v$ equals the source node $u$. For message passing, NBFNet borrows relational operators from KG embeddings, such as translation from TransE, multiplication from DistMult, and rotation from RotatE. Aggregation functions are implemented using permutation-invariant functions from GNN literature, including sum, mean, max, and principal neighborhood aggregation (PNA). Traditionally, edge representations are defined as transition probabilities or lengths. However, since an edge's contribution varies with the query relation, NBFNet parameterizes edge representations as a linear function of the query relation [3]. NBFNet can be interpreted as a novel GNN framework for learning pair representations. Unlike common GNN frameworks, which compute pair representations as two independent node representations $h_q(u)$ and $h_q(u)$, in NBFNet, each node learns a representation conditioned on the source node. The learned pair representation $h_q(u,v)$ is then used to solve the link prediction problem, i.e., predicting the tail entity $v$ given the head entity $u$ and relation $q$. This is formulated as the conditional likelihood of the tail entity $v$ as:

$$p(v|u,q) = \sigma(f(h_q(u,v))) \tag{9}$$

where where $\sigma(\_)$ is the sigmoid function and $f(\_)$ is a feed-forward neural network.

#### 1.2 Gene function prediction task

The following hyperparameters of BioPathNet were optimized for: adversarial temperature (corresponding to the temperature in self-adversarial negative sampling) $\{0.5, 1.0, 2.0\}$; negative samples: $\{32, 64\}$; aggregation function: sum, PNA (Principal Neighborhood Aggregator [4]; number of hidden layers: $\{4, 5, 6\}$; hidden layer dimension: $\{32, 64\}$; learning rate $\{5e-3, 1e-3, 5e-4\}$. Optimal hyperparameter set of BioPathNet {’layers’: 6, ’hidden_dim’: 32, ’num_negatives’: 32, ’lr’: 0.005, ’adversarial_temperature’: 0.5, ’batch_size’: 8, ’epochs’: 8}. Hyperparameters were optimized on the validation MRR.

##### 1.3 Drug repurposing task

For this task, the hyperparameter search for the BioPathNet model considered the parameters dependent $= \{yes, no\}$, number of hidden layers $= \{2, 4, 6, 8\}$, aggregator function $= \{sum, pna\}$, adversarial temperature $= \{0.5, 1, 2, 5\}$, and number of negative samples $= \{32, 64, 128\}$. The best MRR performance in the KG completion validation set determined the final parameter set. The experiments were repeated for five different data split seeds in the TxGNN code. The mean ± standard deviation of performance metrics were reported.

##### 1.4 Synthetic lethality gene pair prediction task

We train BioPathNet for 15 epochs and for five random seeds and tuned hyperparameters such as the number of hidden layers in $\{4, 5\}$, the number of sampled negatives per positive example in $\{32, 64\}$, and the adversarial temperature in $\{0.5, 1.0, 2.0, 5.0\}$. Optimal hyperparameter set of BioPathNet {’layers’: 5, ’hidden_dim’: 32, ’num_negatives’: 64, ’lr’: 0.005, ’adversarial_temperature’: 0.5, ’batch_size’: 64, ’epochs’: 15}.

##### 1.5 LncRNA-target prediction task

The KG constructed by LncTarD 2.0 consists mainly of a densely connected subgraph of 2,646 genes and 6,084 involved regulations, surrounded by many isolated subgraphs containing only two or three genes. Only the largest connected component was kept and split into train, validation, and test set using the python package PyKeen [5], at a ratio of 0.8, 0.1, and 0.1, while ensuring that all nodes were included in the training set with at least one triplet. The optimal hyperparameter set of BioPathNet in this setting was: {’layers’: 6, ’hidden_dim’: 32, ’num_negatives’: 32, ’lr’: 0.005, ’adversarial_temperature’: 0.5, ’batch_size’: 64, ’epochs’: 10}.

##### 1.6 Baseline methods used for comparison

**TransE** The key idea of TransE is to model relationships as translations in the embedding space. For each triplet $(u, r, v)$, the relationship $r$ is modeled as a translation vector such that the embedding of the tail entity $v$ should be close to the embedding of the head entity $u$ plus the embedding of the relationship $r$: $\mathbf{u} + \mathbf{r} \approx \mathbf{v}$. The plausibility of a triplet $(u, r, v)$ is determined by a scoring function based on the distance between $\mathbf{u}+\mathbf{r}$ and $v$: $f(u, r, v) = \|\mathbf{u}+\mathbf{r}-\mathbf{v}\|_2$. TransE is trained to minimize the distance for correct triplets and maximize the distance for incorrect triplets using a margin-based ranking loss [6]. The following hyperparameters were used: {’embedding_dim’: 512, ’num_negatives’: 512, ’lr’: 0.0005, ’epochs’: 100}.

**RotatE** RotatE is a KGE model that represents entities as complex-valued vectors and relationships as rotations in the complex plane. This allows the model to capture various relational patterns, including symmetry, antisymmetry, inversion, and composition. Each entity $u$ and $v$ (head and tail) is represented as a complex vector, and each relationship $r$ is represented as a complex vector with a modulus of 1, representing a rotation in the complex plane. For a triplet $(u, r, v)$, the relationship $r$ rotated $u$ to match $v$, $v \approx u \circ v$, where $\circ$ denotes the element-wise (Hadamard) product. The plausibility of a triplet $(u, r, v)$ is determined by a scoring function based on the distance between $u$ and $v$: $f(u, r, v) = \|\mathbf{u} \circ \mathbf{r} - \mathbf{v}\|_2$. RotatE is trained to minimize the distance for correct triplets and maximize the distance for incorrect triplets using a margin-based ranking loss [7]. The following hyperparameters were used: {’embedding_dim’: 512, ’num_negatives’: 512, ’lr’: 0.0005, ’epochs’: 100}.

**DistMult** DistMult (Multiplicative Distance) is a knowledge graph embedding model that represents entities and relationships as vectors and captures interactions through a multiplicative mechanism. Each entity $u$ and $v$ (head and tail) and each relationship $r$ is represented as a vector in the embedding space. For a triplet $(u, r, v)$, the relationship $r$ interacts multiplicatively with the head entity $u$ to predict the tail entity $v$: $v \approx u \circ r$, where $\circ$ denotes the element-wise (Hadamard) product. The plausibility of a triplet $(u, r, v)$ is determined by a scoring function that measures the similarity between $u \circ r$ and $v$: $f(u, r, v) = -\|\mathbf{u} \circ \mathbf{r} - \mathbf{v}\|_2^2$. DistMult is trained to maximize the score for correct triplets and minimize the score for incorrect triplets using, similarly to TransE, a margin-based ranking loss [8]. The following hyperparameters were used: {'embedding_dim': 512, 'num_negatives': 512, 'lr': 0.001, '$L_3$ regularization' = 1e-4, 'epochs': 100}.

**Relational Graph Convolutional Networks (R-GCNs)** R-GCN is a variant of the Graph Convolutional Network (GCN) specifically designed to handle graphs with multiple types of relations. R-GCN extends GCN to incorporate relational data, making it suitable for KGs. R-GCN employs a neural encoder to learn the representation for each node, while the decoder retains the scoring functions from the KGE models. Initially, the embeddings for each node are initialized [9]. During each layer of the GNN's message passing, messages are passed, aggregated, and used to update the node's embedding. The messages are obtained by applying a $W_{r,M}^{(l)}$ matrix on the embeddings of the previous layer $h_i^{(l-1)}$ separately for each relation $r$, allowing R-GCN to handle graphs with multiple types of edges by learning separate weight matrices for each relation type: $m_{r,i}^{(l)} = W_{r,M}^{(l)} h_i^{(l-1)}$. Next, the incoming messages of each node $v_i$ from the neighboring nodes $N_r(i)$ are aggregated by the taking the average: $\widetilde{m_{r,i}^{(l)}} = \frac{1}{|N_r(l)|} \sum_{j \in N_r(i)} m_{r,j}^{(l)}$. Finally, the node's embedding from the previous layer is updated by $h_i^{(l)} = h_i^{(l-1)} + \sum_{r \in T_R} \widetilde{m_{r,i}^{(l)}}$. This process ensures that each node's embedding is updated with information from its neighbors, considering the type of relation connecting them. The final embeddings of the nodes are derived after $L$ layers of propagation. The hyperparameters for R-GCN were optimized using grid search on the parameters *num_negatives*, *lr* and *hidden_dim*. Two separate searches were conducted: one without the BRG, yielding the best set as {'num_negatives': 64, 'lr': $10^{-4}$, 'layers': 3, 'hidden_dim' = 256}, and another with the BRG, resulting in the best set as {'num_negatives': 32, 'lr': $10^{-5}$, 'layers': 5, 'hidden_dim' = 128}.

**Transductive Graph Neural Network (TxGNN)** TxGNN is a graph neural network-based model designed to predict drug-disease relationships in zero-shot scenarios where there is minimal prior information or treatment history. Utilizing PrimeKG, a comprehensive biomedical knowledge graph, TxGNN employs R-GCNs to learn embeddings of drugs and diseases, capturing complex interactions by mapping them into a shared latent space. This allows TxGNN to predict drug-disease interactions, focusing on indications and contraindications, even for diseases with limited molecular characterization. It achieves this through the use of disease signature vectors, adaptive embedding aggregation, and iterative optimization of non-linear transformations. TxGNN improves drug-disease prediction by splitting training into pre-training and fine-tuning phases. In pre-training, all triplets are used, while fine-tuning focuses on drug-disease relations like `indication`, `contraindications`, and `off-label use`. This is in contrast to BioPathNet, where we do not divide our approach into these two phases or perform pre-training. Instead, we use all triplets containing non-drug-disease relations in the message passing BRG without considering them for supervision. TxGNN was trained using the hyperparameters provided in the original publication: pretraining: {'lr': $1e-3$, 'batch_size': 1024, 'pretrain epochs': 2}; finetuning: {'lr': $5e-4$, 'hidden_dim': 512, 'batch_size': 1024, 'pretrain epochs': 500}. The hyperparameters used in fine-tuning were the same for all disease splits.

**Knowledge Representation for Synthetic Lethality (KR4SL)** KR4SL is a path-representation learning GNN-based method for the prediction of SL gene pairs. The framework begins with the construction of a heterogeneous KG that integrates genes, gene interactions, pathways, GO terms, and an SL graph augmented with reverse and identity edges. This graph is processed through an encoder that identifies relevant relational paths for each primary gene, combining structural graph information with textual semantics of entities. KR4SL utilizes a gated recurrent unit (GRU) to enhance the sequential semantics of the relational paths. As messages propagate through the network, they are enriched by fusing structural and textual information. An attentive aggregation mechanism evaluates the importance of different edges, focusing on the most essential paths for model explanations. Finally, a scoring decoder processes the semantic representations of candidate SL partners, generating scores to rank and select the top-$k$ candidates as the most probable SL partners.

We use the code from GitHub to train KR4SL in the transductive setting. We trained KR4SL on the original (unthresholded) data using the same hyperparameters specified by the authors on GitHub, 'weight decay rate': 0.000089, 'lr': 0.0011, 'batch_size': 50, 'epochs': 15, 'hidden_dim': 48, 'drop_out': 0}. The experiments were run five times with different seeds. The same set of parameters was used for thresholded data.

##### 1.7 Model Evaluation

Models were evaluated by ranking positive triplets $\langle u, r, v\rangle$ of the test set against all negative triplets $\langle u, r, v'\rangle$ that are not present in the KG, following the filtered ranking protocol [8]. Equally, for the reverse relation direction $r^{-1}$, positive triplets of the form $\langle v, r, u\rangle$ are ranked against all negatives $\langle v, r^{-1}, u\rangle$. This approach yields a very stringent evaluation that forces the model to rank positive samples highly. The metrics used are mean rank (MR), mean reciprocal rank (MRR), and hits at k (Hits@$k$). MR is calculated by averaging the ranks $q$ of the positive samples among negatives. Lower values are ideal, with the best value at 1.

$$MR = \frac{1}{|Q|}\sum_{q\in Q} q \tag{10}$$

MRR is the average of the reciprocal rank $\frac{1}{q}$, placing less emphasis on lowly ranked triplets. The values range in $[0, 1]$, and the larger the value, the better the model.

$$MRR = \frac{1}{|Q|}\sum_{q\in Q} \frac{1}{q} \tag{11}$$

Finally, the metric Hits@$k$ provides the probability of correct predictions in the top $k$ predicted triplets made by the model. It can also be considered the percentage of positive triplets in the top $k$ and is, therefore, equivalent to Recall@$k$. Like MRR, values range in $[0, 1]$, and the larger the value, the better the model.

$$H@k = \frac{\left|\{q \in Q : q < k\}\right|}{\left|Q\right|} \tag{12}$$

The best model is selected based on the highest MRR in a validation set.

While the conditional probability is modeled in KGC by predicting the tail entity $v$ given the head entity $u$ and the relation $r$, it might be sensible to evaluate under the joint probability of $u$, $v$, and $r$. Therefore, to remain consistent with the TxGNN we compared against in the drug prediction task, we also used AUPRC, which summarizes the precision and recall at different probability thresholds, as well as specificity and F1 score at the probability threshold of 0.5 (equation 13) using code inherent to TxGNN. They evaluate the performance per disease node.

$$Precision = \frac{TP}{TP + FP},\ Recall = \frac{TP}{TP + FN},\ F1 = 2 \times \frac{Precision \times Recall}{Precision + Recall} \tag{13}$$

TxGNN calculates AUPRC using two strategies. The first compares each positive ground truth item vs. one negative from the list of drugs of a disease area, referred to as `AUPRC 1:1`. The second considers all positive and negative ground truth items, referred to as `AUPRC`. The first metric conveys information on the model's ability to distinguish a positive from a randomly sampled negative in direct comparison. In contrast, the latter focuses on distinguishing positives and negatives across the whole dataset. Furthermore, the `AUPRC` accounts for the positive/negative class imbalance and provides a balanced evaluation of precision and recall. We chose to predominately use the latter, which reflects a real-world scenario of identifying therapeutical opportunities within a given set of drugs.

For the SL prediction task, we compared the seed-wise performance of our model with the performance of KR4SL using metrics inherent to the KR4SL framework's code: NDCG@k, Recall@k, and Precision@k. Moreover, we computed MRR from equation 11 for both BioPathNet and KR4SL by first calculating MRR for each query gene and then averaging gene-wise MRR overall query genes.

Recall@k here is utilized to evaluate how effectively a model predicts SL relationships within its top $k$ predictions and is defined as follows:

$$Recall@k = \frac{1}{N} \sum_{n=1}^{N} \frac{\sum_{i=1}^{k} \mathbb{1}\{G_n^{KR}(i) \in G_n^{KP}\}}{|G_n^{KP}|} \tag{14}$$

where $N$ is the total number of query genes, $G_n^{KP}$ is the set of known SL partner candidates of gene $n$ and $G_n^{KR}$ is the set of top-$k$ SL gene partners for gene $n$ ranked by prediction scores meaning that $G_n^{KR}(i)$ is $i$-th predicted SL partner for gene $n$.

Respectively, Precision@k is employed to assess how effectively a model predicts SL relationships within its top-$k$ predictions, normalized by either the size of the top-$k$ predictions or the size of the known SL relationships, whichever is smaller. Precision@k is defined as follows:

$$Precision@k = \frac{1}{N} \sum_{n=1}^{N} \frac{\sum_{i=1}^{k} \mathbb{1}\{G_n^{KR}(i) \in G_n^{KP}\}}{min\{|G_n^{KP}|, k\}} \tag{15}$$

NDCG@k is used to evaluate a model's capability to prioritize the top-$k$ candidate genes associated with SL relationships and is defined as follows:

$$NDCG@k = \frac{1}{N} \sum_{n=1}^{N} \frac{DCG@k(n)}{IDCG@k(n)} \tag{16}$$

where $DCG@k(n)$ and $IDCG@k(n)$ are defined as follows:

$$DCG@k(n) = \sum_{i=1}^{k} \frac{2^{\mathbb{1}\{G_n^{KR}(i) \in G_n^{KP}\}} - 1}{log_2(i+1)} \tag{17}$$

$$IDCG@k(n) = \sum_{i=1}^{min\{|G_n^{KP}|, k\}} \frac{1}{log_2(i+1)} \tag{18}$$

### 2 Supplementary Tables

#### 2.1 Gene function prediction task

**Supplementary Table 1**: Summary of the dataset for the gene function prediction task: the **number of nodes, edges and relation types** in the background regulatory graph (BRG) obtained from Pathway Commons, as well as train, validation and test set obtained from KEGG.

| | Number of nodes | Number of edges | Number of relations |
|---|---|---|---|
| BRG | 30,885 | 1,884,146 | 13 |
| Train | 7,070 | 22,702 | 1 |
| Valid | 2,348 | 3,243 | 1 |
| Test | 3,533 | 6,487 | 1 |
| Unique count | 31,410 | 1,916,513 | 14 |

**Supplementary Table 2**: Detailed breakdown of the relation types in the functional annotation data set: **Number of edges per relation type** in BRG, train, validation and test set.

| | Relation | Count |
|---|---|---|
| BRG | controls-transport-of-chemical | 3,741 |
| | reacts-with | 4,063 |
| | controls-transport-of | 7,899 |
| | used-to-produce | 14,747 |
| | controls-phosphorylation-of | 17,660 |
| | controls-production-of | 21,262 |
| | consumption-controlled-by | 22,659 |
| | controls-expression-of | 125,860 |
| | catalysis-precedes | 147,948 |
| | in-complex-with | 191,275 |
| | controls-state-change-of | 191,548 |
| | interacts-with | 517,390 |
| | chemical-affects | 618,094 |
| Train | KEGGPathway | 22,702 |
| Valid | KEGGPathway | 3,227 |
| Test | KEGGPathway | 6,438 |

**Supplementary Table 3**: **Best hyperparameter sets for gene function prediction tasks** with and without BRG, *dim* refers to embedding dimension, *L* is the number of hidden layers, *lr* is the learning rate, *Adv. T* the adversarial temperature used in self-adversarial negative sampling, *Aggr. Function* specifies the aggregation function, relevant only for BioPathNet that can choose between *sum* and *PNA*.

| **Model** | *dim* | *L* | *lr* | *NegSam* | *Adv. T* | *Aggr. Function* |
|---|---|---|---|---|---|---|
| TransE w BRG | 128 | - | 1e-3 | 512 | 1 | - |
| TransE w/o BRG | 512 | - | 5e-4 | 512 | 1 | - |
| DistMult w BRG | 128 | - | 1e-3 | 512 | 1 | - |
| DistMult w/o BRG | 512 | - | 1e-3 | 512 | 1 | - |
| RotatE w BRG | 128 | - | 2e-4 | 512 | 1 | - |
| RotatE w/o BRG | 512 | - | 5e-4 | 512 | 1 | - |
| R-GCN w BRG | 128 | 5 | 5e-4 | 32 | 1 | - |
| R-GCN w/o BRG | 256 | 3 | 1e-4 | 64 | 1 | - |
| BioPathNet w BRG | 32 | 6 | 5e-3 | 32 | 1 | PNA |
| BioPathNet w/o BRG | 32 | 6 | 5e-3 | 32 | 1 | PNA |

#### 2.2 Drug repurposing task

**Supplementary Table 4**: **Number of diseases per disease area split**, as well as the number of edges used as BRG and for training, validation, and testing.

| Disease area | Number of diseases | BRG | Train | Valid | Test contraindication | Test indication |
|---|---|---|---|---|---|---|
| Adrenal gland | 6 | 5,728,452 | 33,063 | 4,723 | 303 | 33 |
| Anemia | 19 | 5,705,775 | 33,715 | 4,817 | 752 | 88 |
| Cardiovascular | 111 | 5,695,332 | 30,930 | 4,419 | 4,215 | 453 |
| Cell proliferation | 201 | 5,689,920 | 33,102 | 4,729 | 1,047 | 999 |
| Mental health | 60 | 5,690,512 | 33,443 | 4,778 | 1,567 | 355 |

**Supplementary Table 5**: **Zero-shot prediction scenario for Alzheimer's disease**: diseases included and number of contraindication and indication drugs in custom data split generated following TxGNN code on disease evaluation.

| Disease | ID | # contraindication | # indication |
|---|---|---|---|
| Alzheimer disease | 28,780 | 56 | 8 |
| Alzheimer disease w/o neurofibrillary tangles | 83,960 | 2 | 7 |
| Lewy body dementia | 29,296 | 2 | 0 |
| Pick disease | 28,473 | 27 | 7 |
| Dementia (disease) | 37,573 | 28 | 3 |

**Supplementary Table 6**: **Best Hyperparameter Sets for Each Disease Area Split**, *dim* refers to embedding dimension, *L* is the number of hidden layers, *lr* is the learning rate, *Adv. T* the adversarial temperature used in self-adversarial negative sampling, *Aggr. Function* specifies the aggregation function, in this case PNA was always the best performing.

| **Disease Area** | *dim* | *L* | *lr* | *NegSam* | *Adv. T* | *Aggr. Function* |
|---|---|---|---|---|---|---|
| Adrenal Gland | 32 | 6 | 5e-3 | 64 | 0.5 | PNA |
| Anemia | 32 | 4 | 1e-3 | 64 | 1.0 | PNA |
| Cardiovascular | 32 | 6 | 1e-3 | 64 | 0.5 | PNA |
| Cell Proliferation | 32 | 6 | 1e-3 | 64 | 1.0 | PNA |
| Mental Health | 32 | 4 | 5e-3 | 64 | 1.0 | PNA |
| Alzheimer | 32 | 6 | 5e-3 | 64 | 1.0 | PNA |

**Supplementary Table 7**: **Performances across five different disease area splits**, mean metrics summarized as average across contraindication and indication.

Anemia

| | Recall @20 | AP @20 | MRR @20 | F1 | AUPRC 1:1 | AUPRC |
|---|---|---|---|---|---|---|
| TxGNN | 0.45 | 0.457 | 0.535 | 0.244 | 0.667 | 0.352 |
| BioPathNet | **0.504** | **0.493** | **0.52** | **0.305** | **0.674** | **0.412** |

Adrenal gland

| | Recall @20 | AP @20 | MRR @20 | F1 | AUPRC 1:1 | AUPRC |
|---|---|---|---|---|---|---|
| TxGNN | **0.604** | 0.724 | 0.725 | **0.612** | 0.587 | 0.632 |
| BioPathNet | 0.562 | **0.903** | **0.9** | 0.405 | **0.582** | **0.721** |

Cardiovascular

| | Recall @20 | AP @20 | MRR @20 | F1 | AUPRC 1:1 | AUPRC |
|---|---|---|---|---|---|---|
| TxGNN | 0.1 | 0.17 | 0.199 | 0.059 | 0.606 | 0.108 |
| BioPathNet | **0.166** | **0.23** | **0.249** | **0.074** | **0.605** | **0.136** |

Cell proliferation

| | Recall @20 | AP @20 | MRR @20 | F1 | AUPRC 1:1 | AUPRC |
|---|---|---|---|---|---|---|
| TxGNN | 0.539 | 0.487 | 0.549 | 0.256 | 0.811 | 0.437 |
| BioPathNet | **0.604** | **0.626** | **0.654** | **0.404** | **0.813** | **0.556** |

Mental health

| | Recall @20 | AP @20 | MRR @20 | F1 | AUPRC 1:1 | AUPRC |
|---|---|---|---|---|---|---|
| TxGNN | 0.167 | 0.228 | 0.264 | 0.112 | 0.617 | 0.16 |
| BioPathNet | **0.219** | **0.265** | **0.297** | **0.132** | **0.57** | **0.187** |

#### 2.3 Synthetic lethality gene pair prediction task

**Supplementary Table 8**: **The statistics** of the BRG and SL graph as reported by KR4SL.

| | # Entities | # Relationss | # Triples |
|---|---|---|---|
| SL graph | 9,746 | 1 | 35,374 |
| BRG | 42,547 | 32 | 381,761 |

**Supplementary Table 9**: **Number of edges and source genes** in each data split for various thresholds

| threshold | 0.0 | 0.2 | 0.3 | 0.4 | 0.5 | 0.6 | 0.7 | 0.8 |
|---|---|---|---|---|---|---|---|---|
| # edges (BRG) | 396,619 | 396,221 | 394,922 | 392,429 | 390,151 | 386,705 | 385,090 | 384,900 |
| # genes (BRG) | 14,858 | 14,460 | 13,161 | 10,668 | 8,390 | 4,944 | 3,329 | 3,139 |
| # edges (train) | 9,878 | 9,617 | 8,770 | 7,038 | 5,540 | 3,274 | 2,137 | 2,015 |
| # genes (train) | 5,444 | 5,363 | 4,747 | 3,252 | 2,597 | 1,804 | 846 | 755 |
| # edges (valid) | 3,533 | 3,447 | 3,172 | 2,568 | 2,004 | 1,184 | 764 | 721 |
| # genes (valid) | 2,918 | 2,870 | 2,544 | 1,803 | 1,412 | 946 | 556 | 516 |
| # edges (test) | 7,049 | 6,887 | 6,254 | 5,029 | 3,909 | 2,279 | 1,556 | 1,466 |
| # genes (test) | 4,474 | 4,407 | 3,853 | 2,601 | 2,020 | 1,367 | 739 | 664 |

**Supplementary Table 10**: **Detailed breakdown of the relation types** in the SynLethDB: Number of edges per head and tail type in BRG

| Entity 1 | Relation | Entity 2 | # Triples |
|---|---|---|---|
| gene | participates GpPW | Pathway | 41,726 |
| gene | NOT\|enables | molecular function | 233 |
| gene | NOT\|involved in | biological process | 376 |
| gene | NOT\|is active in | cellular component | 3 |
| gene | NOT\|located in | cellular component | 109 |
| gene | NOT\|part of | cellular component | 134 |
| gene | is active in | cellular component | 8,749 |
| gene | located in | cellular component | 51,382 |
| gene | part of | cellular component | 60,679 |
| gene | acts upstream of or within | biological process | 202 |
| gene | acts upstream of | biological process | 158 |
| gene | acts upstream of negative effect | biological process | 2 |
| gene | acts upstream of or within negative effect | biological process | 2 |
| gene | acts upstream of or within positive effect | biological process | 7 |
| gene | acts upstream of positive effect | biological process | 15 |
| gene | colocalizes with | cellular component | 874 |
| gene | contributes to | molecular function | 752 |
| gene | enables | molecular function | 54,511 |
| gene | involved in | biological process | 105,623 |
| gene | NOT\|colocalizes with | cellular component | 9 |
| gene | NOT\|contributes to | molecular function | 3 |
| gene | NOT\|acts upstream of or within | biological process | 1 |
| gene | NOT\|acts upstream of or within negative effect | biological process | 1 |
| biological process | happens during | biological process | 7 |
| biological process | has part | biological process | 144 |
| biological process | has part | molecular function | 94 |
| biological process | is a | biological process | 33158 |
| biological process | negatively regulates | biological process | 1,858 |
| biological process | negatively regulates | molecular function | 192 |
| biological process | occurs in | cellular component | 119 |
| biological process | part of | biological process | 3,144 |
| biological process | positively regulates | biological process | 1,834 |
| biological process | positively regulates | molecular function | 200 |
| biological process | regulates | biological process | 2,092 |
| biological process | regulates | molecular function | 218 |
| cellular component | has part | cellular component | 154 |
| cellular component | is a | cellular component | 3,224 |
| cellular component | part of | cellular component | 1,222 |
| molecular function | has part | molecular function | 167 |
| molecular function | is a | molecular function | 7,493 |
| molecular function | negatively regulates | molecular function | 50 |
| molecular function | occurs in | cellular component | 31 |
| molecular function | part of | biological process | 722 |
| molecular function | part of | molecular function | 8 |
| molecular function | part of | Pathway | 1 |
| molecular function | positively regulates | molecular function | 40 |
| molecular function | regulates | biological process | 1 |
| molecular function | regulates | molecular function | 36 |

**Supplementary Table 11**: **Best hyperparameter set for Synthetic lethality gene pair prediction task**, *dim* refers to embedding dimension, *L* is the number of hidden layers, *lr* is the learning rate, *Adv. T* the adversarial temperature used in self-adversarial negative sampling, *Aggr. Function* specifies the aggregation function.

| **Model** | *dim* | *L* | *lr* | *NegSam* | *Adv. T* | *Aggr. Function* |
|---|---|---|---|---|---|---|
| BioPathNet | 32 | 5 | 5e-3 | 64 | 0.5 | PNA |

**Supplementary Table 12**: **Performance comparison between BioPathNet and state-of-the-art** SL gene pair prediction method KR4SL for **unthresholded data**.

| | seed 1234 | | seed 1235 | | seed 1236 | | seed 1237 | | seed 1238 | |
|---|---|---|---|---|---|---|---|---|---|---|
| metric | KR4SL | BioPathNet | KR4SL | BioPathNet | KR4SL | BioPathNet | KR4SL | BioPathNet | KR4SL | BioPathNet |
| MRR | 0.285 | 0.295 | 0.284 | 0.300 | 0.285 | 0.297 | 0.280 | 0.294 | 0.284 | 0.299 |
| NDCG@10 | 0.341 | 0.364 | 0.341 | 0.362 | 0.343 | 0.360 | 0.337 | 0.366 | 0.340 | 0.366 |
| NDCG@20 | 0.356 | 0.383 | 0.355 | 0.381 | 0.357 | 0.378 | 0.352 | 0.383 | 0.354 | 0.384 |
| NDCG@50 | 0.368 | 0.400 | 0.368 | 0.397 | 0.369 | 0.394 | 0.364 | 0.400 | 0.367 | 0.401 |
| Precision@10 | 0.434 | 0.451 | 0.439 | 0.455 | 0.437 | 0.450 | 0.435 | 0.452 | 0.437 | 0.455 |
| Precision@20 | 0.485 | 0.515 | 0.484 | 0.519 | 0.486 | 0.511 | 0.484 | 0.511 | 0.485 | 0.516 |
| Precision@50 | 0.529 | 0.580 | 0.533 | 0.581 | 0.532 | 0.572 | 0.530 | 0.574 | 0.533 | 0.580 |
| Recall@10 | 0.424 | 0.440 | 0.428 | 0.444 | 0.427 | 0.440 | 0.424 | 0.441 | 0.426 | 0.444 |
| Recall@20 | 0.480 | 0.511 | 0.480 | 0.515 | 0.482 | 0.507 | 0.479 | 0.507 | 0.481 | 0.512 |
| Recall@50 | 0.528 | 0.579 | 0.532 | 0.580 | 0.531 | 0.571 | 0.529 | 0.573 | 0.532 | 0.579 |

**Supplementary Table 13**: **Performance comparison between BioPathNet and state-of-the-art** SL gene pair prediction method KR4SL for **thresholded data**.

| | seed 1234 | | seed 1235 | | seed 1236 | | seed 1237 | | seed 1238 | |
|---|---|---|---|---|---|---|---|---|---|---|
| metric | KR4SL | BioPathNet | KR4SL | BioPathNet | KR4SL | BioPathNet | KR4SL | BioPathNet | KR4SL | BioPathNet |
| MRR | 0.338 | 0.360 | 0.339 | 0.360 | 0.336 | 0.348 | 0.333 | 0.362 | 0.339 | 0.353 |
| NDCG@10 | 0.398 | 0.420 | 0.401 | 0.416 | 0.399 | 0.406 | 0.396 | 0.418 | 0.401 | 0.412 |
| NDCG@20 | 0.416 | 0.437 | 0.415 | 0.434 | 0.415 | 0.427 | 0.412 | 0.436 | 0.416 | 0.432 |
| NDCG@50 | 0.428 | 0.453 | 0.428 | 0.451 | 0.428 | 0.443 | 0.425 | 0.452 | 0.429 | 0.448 |
| Precision@10 | 0.507 | 0.525 | 0.512 | 0.519 | 0.513 | 0.508 | 0.511 | 0.519 | 0.513 | 0.517 |
| Precision@20 | 0.569 | 0.583 | 0.560 | 0.581 | 0.570 | 0.579 | 0.566 | 0.581 | 0.565 | 0.584 |
| Precision@50 | 0.611 | 0.641 | 0.608 | 0.644 | 0.615 | 0.641 | 0.613 | 0.638 | 0.612 | 0.642 |
| Recall@10 | 0.496 | 0.514 | 0.501 | 0.508 | 0.501 | 0.498 | 0.499 | 0.507 | 0.501 | 0.506 |
| Recall@20 | 0.564 | 0.579 | 0.556 | 0.577 | 0.566 | 0.575 | 0.562 | 0.576 | 0.561 | 0.580 |
| Recall@50 | 0.610 | 0.640 | 0.608 | 0.643 | 0.614 | 0.640 | 0.612 | 0.637 | 0.611 | 0.641 |

#### 2.4 LncRNA-gene target prediction task

**Supplementary Table 14**: Summary of the functional annotation data set: **number of nodes, edges and relation types** in the background regulatory graph (BRG), as well as train, validation and test sets.

| | Number of nodes | Number of edges | Number of relations |
|---|---|---|---|
| BRG | 15,806 | 1,035,133 | 7 |
| Train | 2,646 | 4,867 | 6 |
| Valid | 533 | 608 | 6 |
| Test | 535 | 609 | 6 |
| Unique count | 16,844 | 1,041,217 | 13 |

**Supplementary Table 15**: Detailed breakdown of the relation types in the functional annotation data set: **Number of edges per relation type** in BRG, train, validation and test set.

| | Relation | Count |
|---|---|---|
| BRG | interacts-with | 469,203 |
| | in-complex-with | 154,868 |
| | controls-state-change-of | 144,259 |
| | catalysis-precedes | 141,554 |
| | controls-expression-of | 114,698 |
| | controls-phosphorylation-of | 7,246 |
| | controls-transport-of | 3,305 |
| Train | ceRNA or sponge | 2,305 |
| | expression association | 1,530 |
| | interact with protein | 366 |
| | transcriptional regulation | 351 |
| | epigenetic regulation | 272 |
| | interact with mRNA | 43 |
| Valid | ceRNA or sponge | 275 |
| | expression association | 208 |
| | interact with protein | 48 |
| | transcriptional regulation | 44 |
| | epigenetic regulation | 31 |
| | interact with mRNA | 2 |
| Test | ceRNA or sponge | 247 |
| | expression association | 228 |
| | interact with protein | 52 |
| | transcriptional regulation | 47 |
| | epigenetic regulation | 28 |
| | interact with mRNA | 7 |

**Supplementary Table 16**: **Best hyperparameter set for LncRNA-gene target prediction task**, *dim* refers to embedding dimension, *L* is the number of hidden layers, *lr* is the learning rate, *Adv. T* the adversarial temperature used in self-adversarial negative sampling, *Aggr. Function* specifies the aggregation function.

| **Model** | *dim* | *L* | *lr* | *NegSam* | *Adv. T* | *Aggr. Function* |
|---|---|---|---|---|---|---|
| BioPathNet | 32 | 6 | 5e-3 | 64 | 0.5 | PNA |

### 3 Supplementary Figures

#### 3.1 Supplementary Figure 1

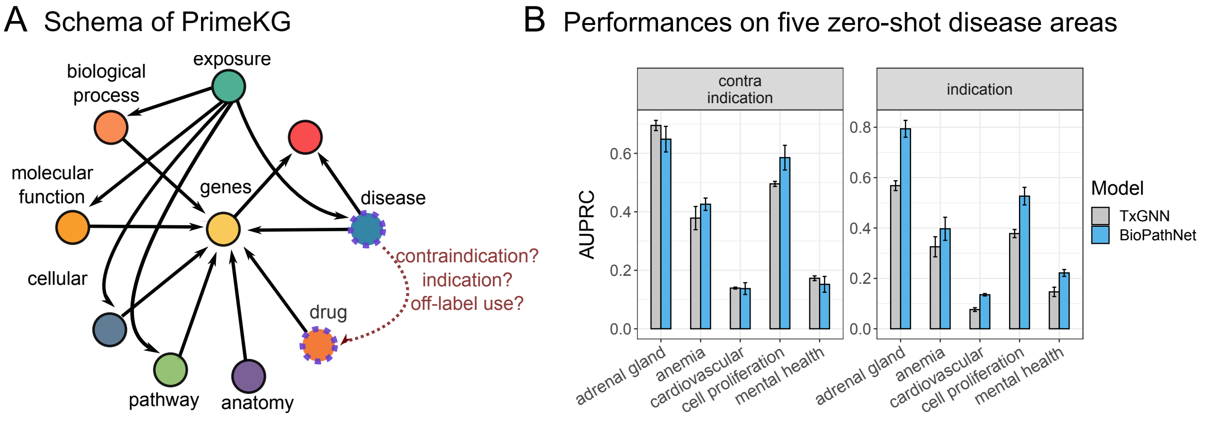

**Supplementary Figure 1**: **BioPathNet's performance on PrimeKG A**) PrimeKG schema: a multi-modal knowledge graph with 10 biological node types (e.g., protein, disease, drug) and over 5 million relations, for predicting drug-disease interactions. **B**) AUPRC performance in zero-shot prediction across five disease area splits (adrenal gland, anemia, cardiovascular, cell proliferation, mental health), calculated by comparing ground truth positives and negatives, then averaging across all diseases in each area.

#### 3.2 Supplementary Figure 2

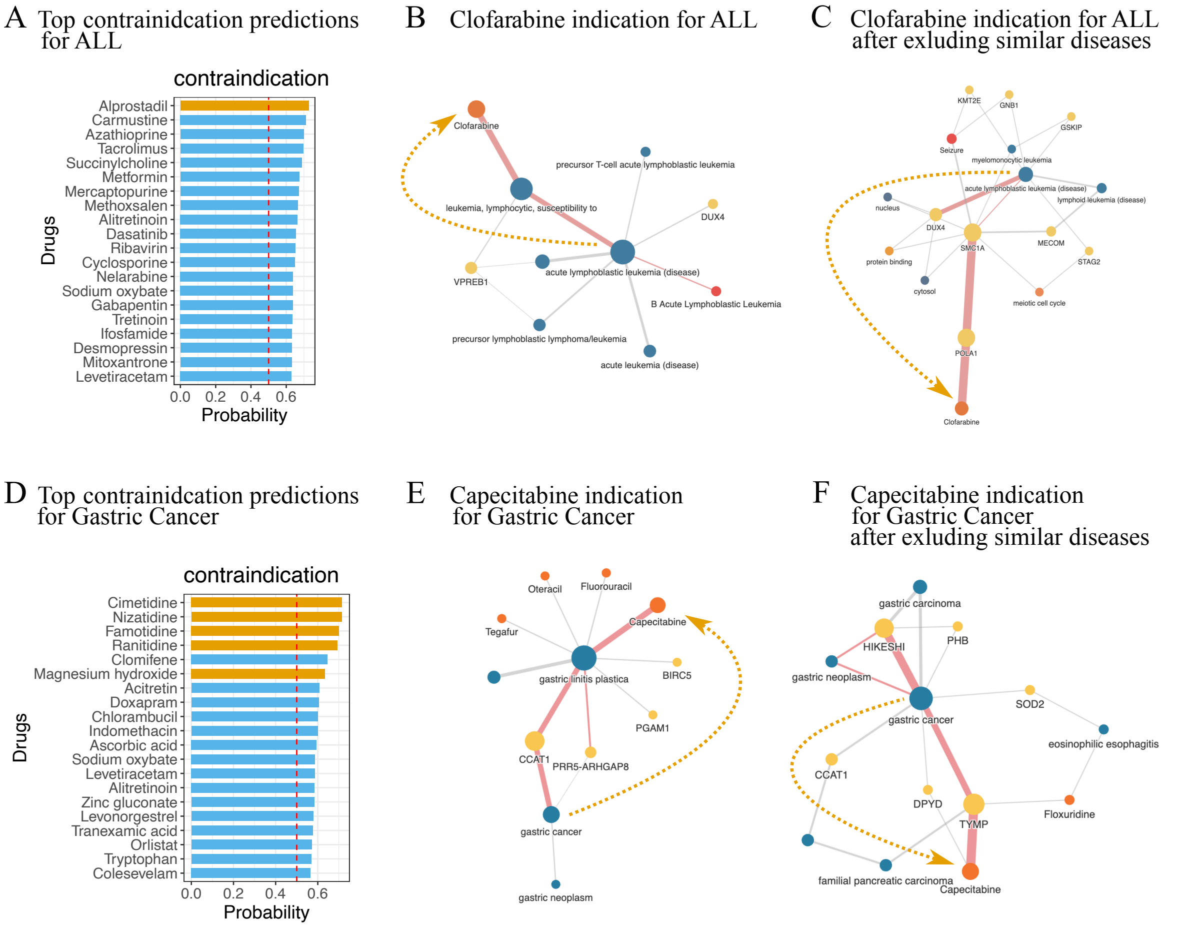

**Supplementary Figure 2**: **Inspecting BioPathNet predictions in zero-shot learning A**) Top 20 predictions of contraindication for ALL. Path interpretation for the prediction of Clorafabine as an indication for ALL in near-shot learning scenario (**B**) and in zero-shot scenario (**C**). **D**) Top 20 predictions of contraindication for Gastric Cancer. Path interpretation for the prediction of Capecitabine as an indication for Gastric Cancer in near-shot learning scenario (**E**) and in zero-shot scenario (**F**). Known contraindications, included in the ground truth of PrimeKG, are highlighted in orange, while while newly predicted contraindications are in light blue. The visualization (B-C and E-F) shows the top 10 significant paths used by BioPathNet for prediction, with edge widths representing weights and the highest-weight path highlighted in red.

#### 3.3 Supplementary Figure 3

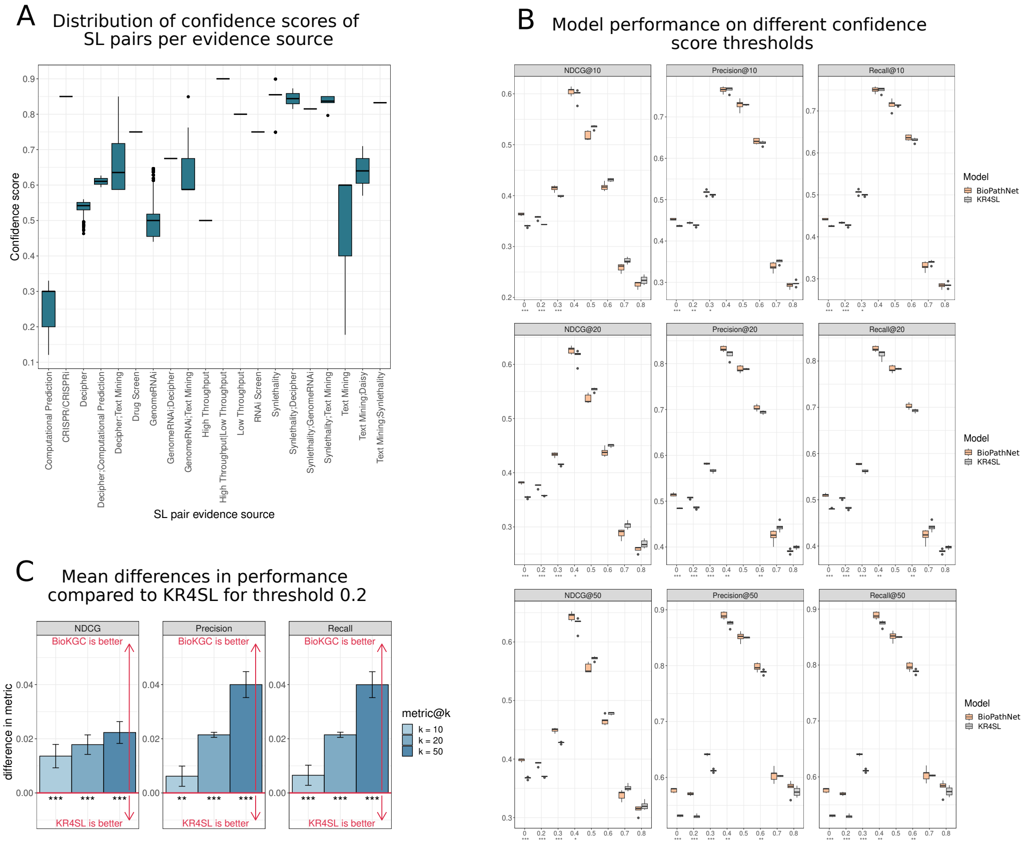

**Supplementary Figure 3**: **Comparison of BioPathNet with state-of-the-art SL gene pair prediction algorithm KR4SL for varying SL confidence thresholds: A**) Distribution of confidence scores of SL pairs per evidence source as provided by SynLethDB-v2.0 as the "r.statistic_score". **B**) Performance comparison of BioPathNet and KR4SL with performance metrics reported across thresholds applied to the confidence score of SL evidence. **C**) Mean difference in performances between BioPathNet and KR4SL given as NDCG@k, Precision@k, and Recall@k, $k \in \{10, 20, 50\}$, for both methods trained on SL pairs which were filtered to have a confidence score of at least 0.2.

#### 3.4 Supplementary Figure 4

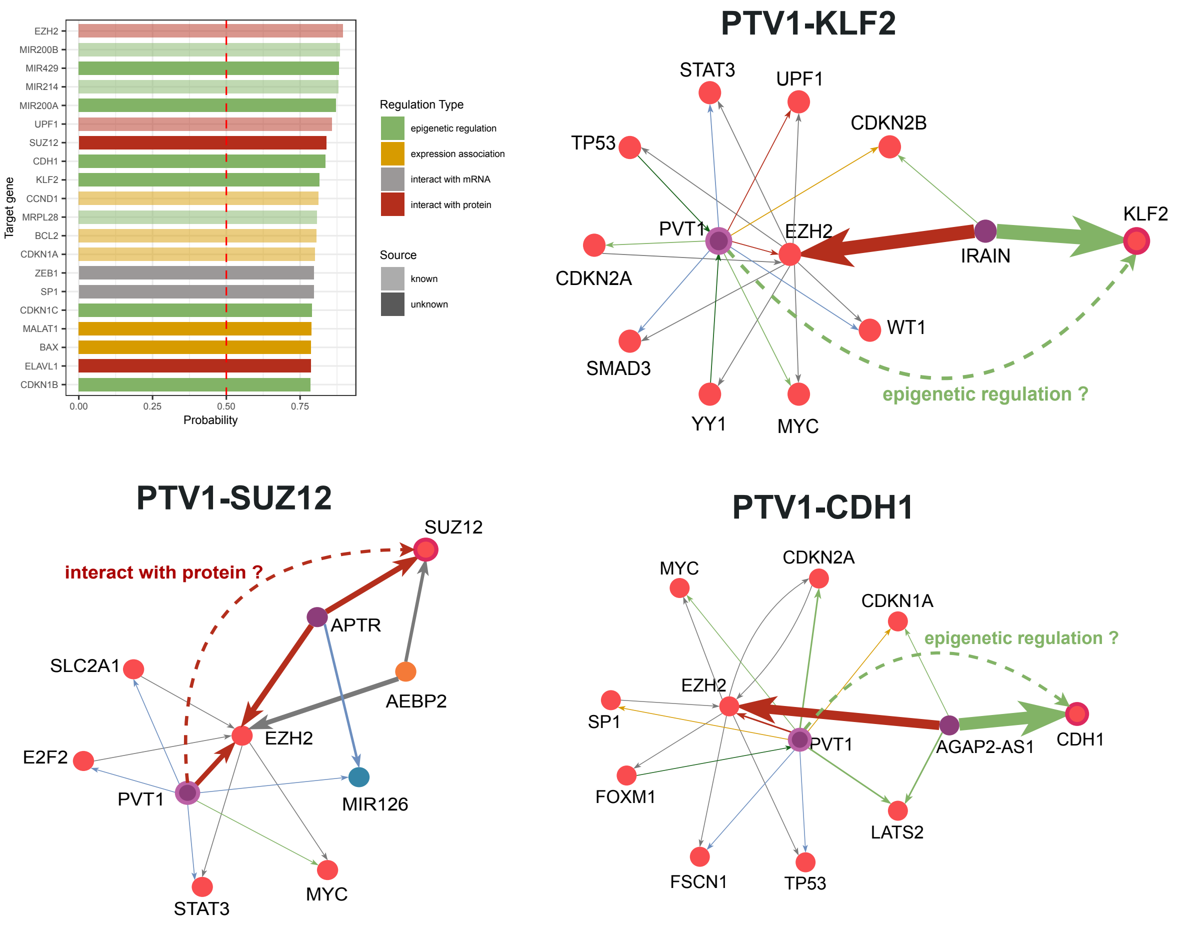

**Supplementary Figure 4**: **Prediction of novel incRNA-target regulatory interactions. A** presents BioPathNet predicted targets for the cancer lncRNA PVT1, ranked by prediction probability, similar to Figure 6A. Additionally, regulation types are distinguished using different colors: green for epigenetic regulation, yellow for expression association, gray for interaction with mRNA, and red for interaction with protein. Known triples are shown with 50% transparency, while novel triples are displayed with full opacity. **B-D** Explanations for top 3-5 novel predicted targets of PVT1: SUZ12, CDH1, and KLF2.
